# Mechanistic and Epigenetic Partitioning of Lamina-Associated Chromatin Revealed by a Genome-Wide Imaging Screen

**DOI:** 10.1101/2025.08.13.670143

**Authors:** Patrick J. Walsh, Elizabeth B. Kraeutler, Ricardo Linares-Saldana, May Wai, Son C. Nguyen, Shuo Zhang, Parisha P. Shah, Daniel S. Park, Haris A. Muzaffar, Rajan Jain, Eric F. Joyce

**Author notes:** Correspondence (EJ) and (RJ). Contributed equally (Patrick J. Walsh, Elizabeth B. Kraeutler).

## Abstract

The nuclear periphery is a key site for heterochromatin organization in eukaryotic cells, where lamina-associated domains (LADs) promote transcriptional repression and genome stability. Despite their importance, the mechanisms governing LAD positioning in human cells remain poorly understood. To this end, we performed a genome-wide imaging-based siRNA screen and identified over 100 genes critical for perinuclear LAD localization, with a striking enrichment for RNA-binding proteins. Among these, hnRNPK emerged as a key regulator, required for the perinuclear positioning of approximately two-thirds of LADs genome-wide. Loss of hnRNPK led to LAD repositioning away from the nuclear periphery without altering their heterochromatin state, yet resulted in misexpression of genes within these domains. Notably, hnRNPK-sensitive LADs are uniquely marked by both H3K9me2 and H3K27me3, distinguishing them from hnRNPK-insensitive LADs that are enriched for H3K9me2 and H3K9me3. These findings reveal at least two mechanistically and epigenetically distinct LAD classes, suggesting that specialized pathways underlie their spatial organization. Our results uncover a pivotal role for hnRNPK in regulating the spatial organization of chromatin and highlight the broader diversity of LAD localization mechanisms.

## Introduction

The nuclear lamina plays a crucial role in the spatial organization of the genome in eukaryotic cells, serving as a scaffold for transcriptionally repressive heterochromatin^1–3^. Lamina-associated domains (LADs), which constitute approximately 30-40% of the genome, are key to this organization^4,5^. These domains are typically gene-poor, transcriptionally silent, late-replicating, and enriched for repressive histone modifications, prominently H3K9me2, H3K9me3, and H3K27me3^6–15^. This conserved organization has been implicated in various biological processes, including development, differentiation, and genome stability^5,7,16–27^, and its disruption has been linked to aging^28–31^ and human disease^32–37^.

Since their initial description, multiple studies have revealed that LADs are not a uniform entity but instead comprise distinct subsets with varied molecular, genetic, and epigenetic features^7,8,11–14,38,39^. These differences include variation in repeat content, histone modifications, and dynamic behavior across cell types. Such heterogeneity suggests that distinct molecular mechanisms may underlie the spatial positioning of specific LAD subsets. Supporting this, a recent study demonstrated that deletion of four independent lamina components was required to disrupt LAD organization, pointing to the presence of multiple, parallel tethering mechanisms^26^. Several lines of evidence also suggest that non-lamina proteins can contribute to gene and/or LAD positioning^16,24,25,40–46^. However, many of these factors either lack clear orthologs in humans or specific roles in heterochromatin spatial positioning and LAD organization. A comprehensive identification of such factors in human cells is therefore essential to fully define the molecular networks that govern chromatin-lamina interactions.

To address this gap, we leveraged our high-throughput DNA/RNA labeling with the Oligopaints (HiDRO) platform^47^ to perform an unbiased genome-wide screen for regulators of LAD positioning in human cells. This screen uncovered over 100 candidate regulators, including a striking enrichment of RNA-processing proteins, and identified hnRNPK as a key organizer of peripheral heterochromatin. Together, these findings reveal that distinct subsets of LADs rely on different molecular mechanisms for their spatial positioning, underscoring the functional and mechanistic heterogeneity within the lamina-associated genome.

## Results

### A genome-wide HiDRO screen for genes important for the spatial positioning of heterochromatin at the nuclear periphery

To identify novel regulators of LAD positioning, we first mapped perinuclear chromatin genome-wide in HCT116 cells using Lamin B1 ChIP-seq. LADs were called using a Hidden Markov Model (HMM) and, consistent with other cell types and prior reports, were on average ∼1 Mb in length and collectively spanned ∼30% of the genome (Fig. 1a). To validate the LAD map, we performed ChIP-seq to the perinuclear heterochromatin-associated histone post-translational modification H3K9me2 and observed strong genome-wide correlation with Lamin B1 enrichment, consistent with previous findings^6,8,16,48,49^ (Supp. Fig. 1a). We then designed Oligopaint DNA FISH probes (Supplementary Table 1) targeting a prominent LAD on chromosome 2q and a nonLAD region on the more centrally located chromosome 22q (Fig. 1b). Using our high-throughput DNA/RNA labeling with Oligopaints (HiDRO) platform^47^ and automated image analysis, we quantified the peripheral distance of each allele, from the centroid of each FISH signal to the nearest nuclear edge as defined by DAPI staining. Across seven replicates each, LAD alleles localized significantly closer to the periphery than nonLAD alleles (Fig. 1c): ∼46% of LAD alleles resided within 1 µm of the nuclear periphery, a zone previously suggested to constrain LADs during interphase (Kind et al. 2013), compared to only ∼8% of nonLAD alleles (Supp. Fig. 1b).

**Figure 1.**
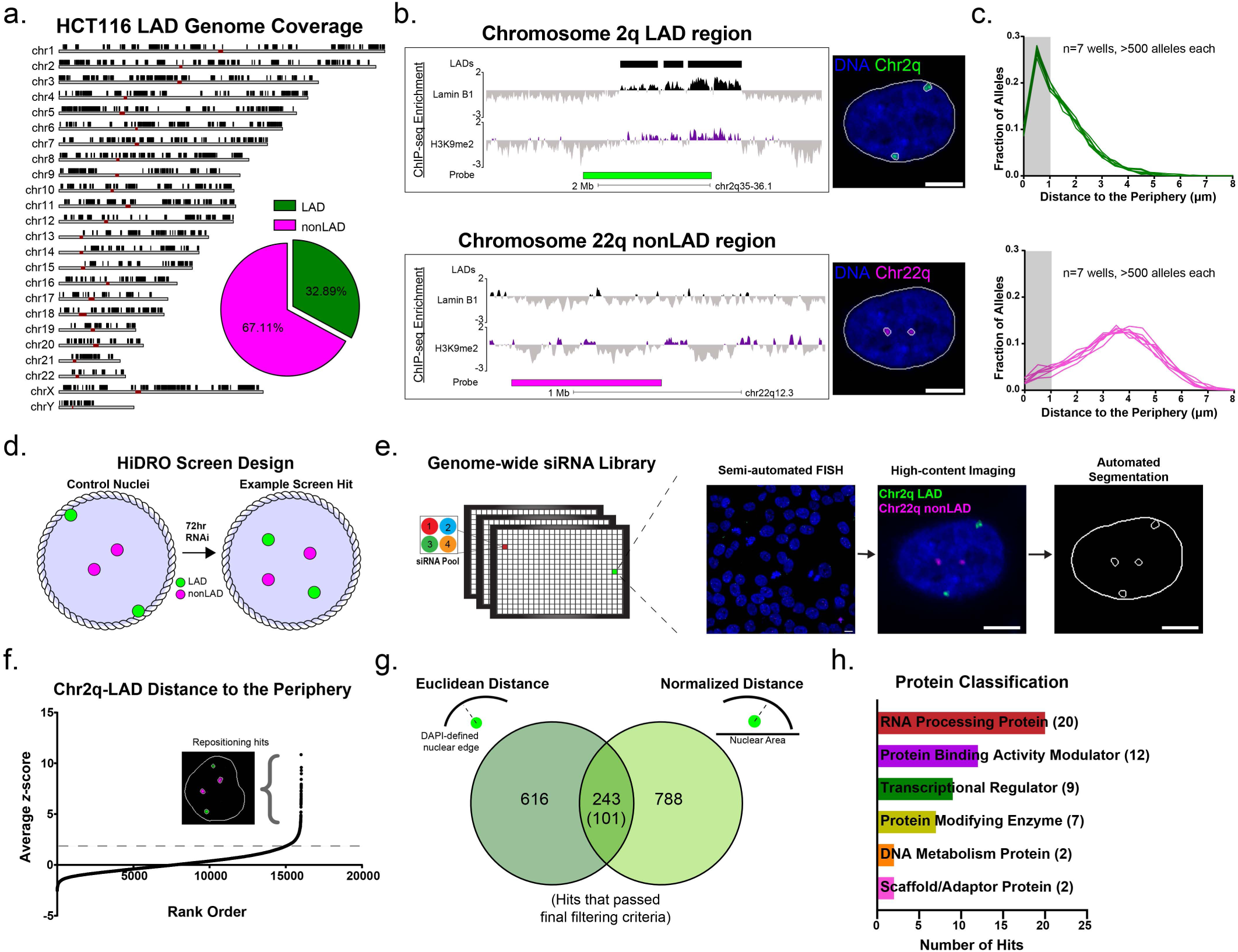
HiDRO Screen for regulators of LAD spatial positioning. a) Ideogram displaying chromosomal distribution of LADs (black) mapped by Lamin B1 ChIP-Seq in HCT116 cells. Maroon bands represent centromeres. Inset, proportion of the hg38 genome (2.8 billion bp) covered by LADs (green) vs nonLAD (magenta). Coverage is the percent of base pairs within LADs. b) Top, Lamin B1 (black) and H3K9me2 (purple) ChIP-seq tracks shown with LAD calls shown in black above. Oligopaint DNA-FISH probe (indicated in green) targeting a LAD on chromosome 2 (chr2q35-36.1). Representative DNA-HiDRO image of Chr2q-LAD in control nucleus (green). The solid white line indicates the nuclear edge determined by DAPI staining (blue). Scale bar, 5 μm. [Bottom] Lamin B1 (black) and H3K9me2 (purple) ChIP-seq tracks shown. Oligopaint DNA-FISH probe (indicated in magenta) targeting a nonLAD region on chromosome 22 (chr22q12.3). Representative DNA-HiDRO image of Chr22q-nonLAD (magenta) in control nucleus. The solid white line indicates the nuclear edge determined by DAPI staining (blue). Scale bar, 5 μm. c) Distribution of distance to the periphery for the Chr2q-LAD (Top, green, n=7 wells) and the Chr22q-nonLAD (Bottom, magenta, n=7 wells). Shaded gray region highlights the fraction of alleles within 1 μm of the nuclear periphery. d) Schematic of example nuclei showing repositioning of the Chr2q-LAD (green) away from the nuclear periphery following RNAi. e) Workflow of HiDRO screen for regulators of LAD positioning. Scale bars, 10 μm (field, left), 5 μm (nucleus, middle), 5 μm (segmentation, right). f) Rank-ordered robust *z*-scores for the Euclidean distance of the Chr2q-LAD to the nuclear periphery for all genes passing the cell count threshold. *Z*-scores are the average of two biological replicates. Inset is an example nucleus with the Chr2q-LAD repositioned away from the nuclear periphery. g) Venn diagram of genes that significantly increased the Chr2q-LAD Euclidean and normalized distance to the nuclear periphery. n = 1,647 genes total. Inset schematics depict how distance measurements were determined. h) Protein classes of primary screen hits that were selected for validation. Protein classes for 52 of 101 primary hits are shown.

We next scaled our HiDRO technology to perform a genome-wide siRNA screen for factors required for the perinuclear localization of the chr2q LAD (Fig. 1d). HCT116 cells were seeded onto 18,218 pre-aliquoted pools of siRNAs in 384-well plates to induce knockdown (KD) of targets for 72 hours. Following fixation, semi-automated Oligopaint FISH was performed to label the chr2q LAD and chr22q nonLAD regions, and plates were imaged on a high-content confocal microscope (Fig. 1e). The primary screen, conducted in duplicate, comprised 40,656 FISH assays and yielded measurements from over 20 million nuclei. Hits were identified based on robust *z*-scores for the Euclidean distance between the chr2q LAD and the nuclear (DAPI) edge calculated from a minimum of 100 nuclei per replicate (Fig. 1f, Supplementary Table 2). Replicates showed strong correlation (Spearman r=0.79), supporting the reproducibility of the screen (Supp. Fig. 1c). Approximately 12% of KDs yielded insufficient cell counts (<100 nuclei per replicate) and were therefore excluded from further analysis (Supp. Fig. 1d). To account for changes in nuclear size that could confound interpretation of distance measurements, *z*-scores were also calculated for distances following normalization to the nuclear area for each nucleus (Supp. Fig. 1e). This analysis yielded 243 siRNAs that increased the distance from the periphery across both replicates and regardless of whether normalized or non-normalized *z*-scores were used (Fig. 1g). However, to further refine this hit list, we excluded knockdowns that (i) increased the distance of the chr22q nonLAD probe to the nuclear edge or (ii) targeted genes not expressed in HCT116 cells. Applying these filtering criteria, we isolated 101 hits (Supplementary Table 3). Gene ontology analysis^50^ revealed diverse molecular functions among the 101 genes, with nearly half (48%) grouped into four major categories: RNA processing proteins, protein-modifying enzymes, gene-specific transcriptional regulators, and protein-binding activity modulators (Fig. 1h, Supp. Fig. 1f). Notably, RNA processing factors were unexpectedly enriched, comprising ∼20% of hits.

### Validation and functional prioritization of hits identify hnRNPK

To prioritize hits from the primary screen for follow-up analysis, we selected 52 candidate genes for validation, representing six protein classes with potential mechanistic relevance to nuclear lamina-associated chromatin organization (Fig. 1h). Each gene was individually targeted with four non-overlapping siRNA duplexes in triplicate wells, totaling 624 FISH reactions excluding controls (Fig. 2a, left). Using HiDRO to relabel the same chr2q LAD as in the primary screen, we observed that 51 of 52 candidates (98%) reproduced the repositioning phenotype with at least one duplex (Fig. 2a, right; Supplementary Table 4). Importantly, 42 of these genes were validated with two or more duplexes, representing our highest-confidence hits.

**Figure 2.**
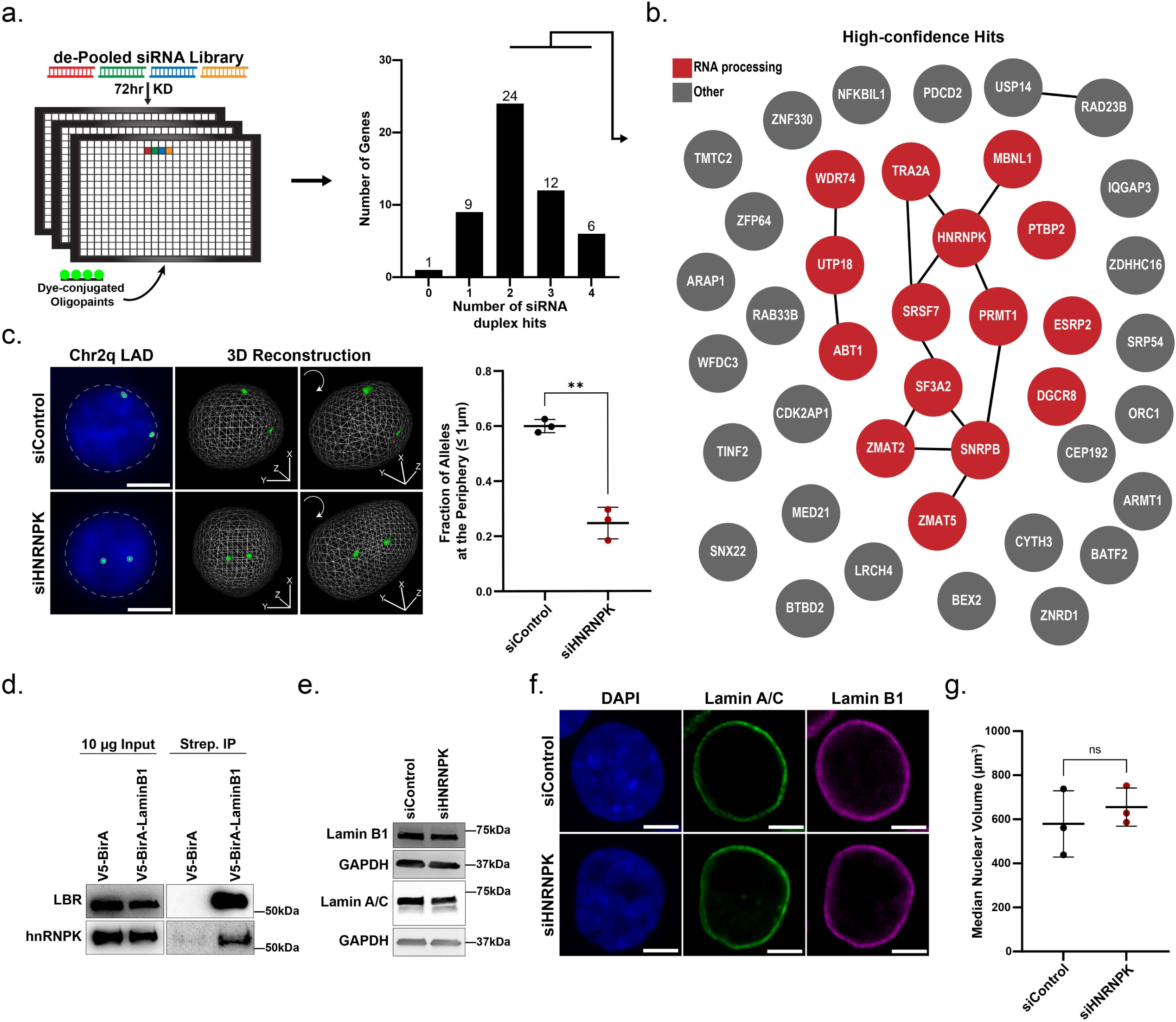
Validation and functional prioritization of hits identify hnRNPK. a) (Left) Validation HiDRO screen of selected primary hits with four non-overlapping siRNA duplexes in separate wells, then applied DNA FISH to the Chr2q-LAD. Each duplex was tested in triplicate. (Right) The number of genes that replicated an increased distance phenotype for the Chr2q-LAD for the number of individual siRNA duplexes. b) STRING network analysis of primary screen hits with at least two validated duplexes. Red bubbles designate genes identified as being related to RNA processing. c) (Left) Representative DNA FISH image of the Chr2q-LAD (green) in control and *HNRNPK* KD nuclei with corresponding 3D image reconstruction. Dashed white line indicates nuclear edge as defined by DAPI staining (blue). Scale bar, 5 μm. [Right] Quantification of 3D DNA FISH analyses showing the fraction of Chr2q-LAD alleles within 1 μm of the nuclear periphery in control (black) and *HNRNPK* KD (dark red). Each dot represents an individual biological replicate, n ≥ 300 nuclei per replicate. Error bars represent mean and SD of replicates. ** *P* = 0.0035, Unpaired t test with Welch’s correction. d) Western blot for Lamin B receptor (LBR) and hnRNPK in HCT116 cells after Turbo-ID biotin labeling from input (left) and streptavidin pull-down (right) using chromatin fractionations. e) Western blot showing levels of Lamin B1 and Lamin A/C in siControl and *HNRNPK* KD whole cell lysates. GAPDH is a loading control. f) Immunofluorescence for Lamin A/C (green) and Lamin B1 (magenta) in siControl and *HNRNPK* KD nuclei. DAPI staining (blue) to label DNA. Scale bars, 5 μm. g) Quantification of nuclear volumes from siControl (black) and *HNRNPK* KD (dark red) nuclei (from 3D DNA FISH experiments in Fig. 2c). Each dot represents the median nuclear volume of an individual biological replicate, n = ≥ 300 nuclei per replicate. Not significant (ns) *P =* 0.50, Unpaired t test with Welch’s correction.

We next performed network analysis^51^, which revealed a prominent enrichment of RNA processing factors (15 of the 42 hits), nine of which formed a single, interconnected subnetwork (Fig. 2b). Among these, the RNA-binding protein hnRNPK emerged as a major node in the network. It was one of only six genes to validate across all four siRNA duplexes (Fig. 2a, right; Supplementary Table 4), and the only factor in the network previously associated with a nuclear positioning phenotype in human cells^41^. No other factors in the hnRNP family were identified in the screen, suggesting a robust and unique role in LAD organization. We confirmed efficient depletion of hnRNPK following 72-hour siRNA treatment by western blot (Supp. Fig. 2a-b). Using a 3D FISH protocol, we also validated the repositioning of the chr2q LAD away from the nuclear periphery across multiple replicates (Fig. 2c, Supp. Fig. 2c). This phenotype was also observed in BJ fibroblasts, suggesting that the role of hnRNPK in LAD positioning is conserved in a non-transformed cell type (Supp. Fig. 2d).

Interestingly, numerous studies have linked hnRNPK to multiple forms of heterochromatin, and large-scale proteomic analyses have identified interactions with nuclear lamina components^13,52–58^. We confirmed hnRNPK association with the nuclear lamina in HCT116 cells using a Lamin B1–BirA fusion protein to perform Turbo-ID proximity labeling^59^. Pull-downs from chromatin fractions revealed enrichment of known lamina-associated proteins, including Lamin B receptor (LBR), in addition to hnRNPK, in Lamin B1-BirA samples compared to the BirA-alone control (Fig. 2d, Supp. Fig. 2e-f). To test whether hnRNPK loss impacts nuclear lamina components, we assessed Lamin B1 and Lamin A/C expression and localization following hnRNPK depletion. Total protein levels of both lamina factors remained unchanged, and both continued to localize to the nuclear periphery, indicating that the lamina itself remains intact (Fig. 2e-f, Supp. Fig. 2g). Nuclear size was also unchanged upon hnRNPK depletion (Fig. 2g). Altogether, these results show that hnRNPK is required for the perinuclear positioning of the chr2q LAD without altering lamina expression or localization.

### hnRNPK regulates LAD organization genome-wide

To determine whether hnRNPK broadly regulates the perinuclear positioning of heterochromatin, we performed Lamin B1 ChIP-seq in HCT116 cells following hnRNPK depletion (Fig. 3a). Highly concordant biological replicates (n = 3 for each condition) were merged for analysis (Supp. Fig. 3a) and LADs were identified using a Hidden Markov Model (HMM). Consistent with our FISH results, hnRNPK KD led to a marked reduction in Lamin B1 enrichment at the chr2q LAD locus with a concomitant loss of any LAD designation in this region (Fig. 3a, red arrow).

**Figure 3:**
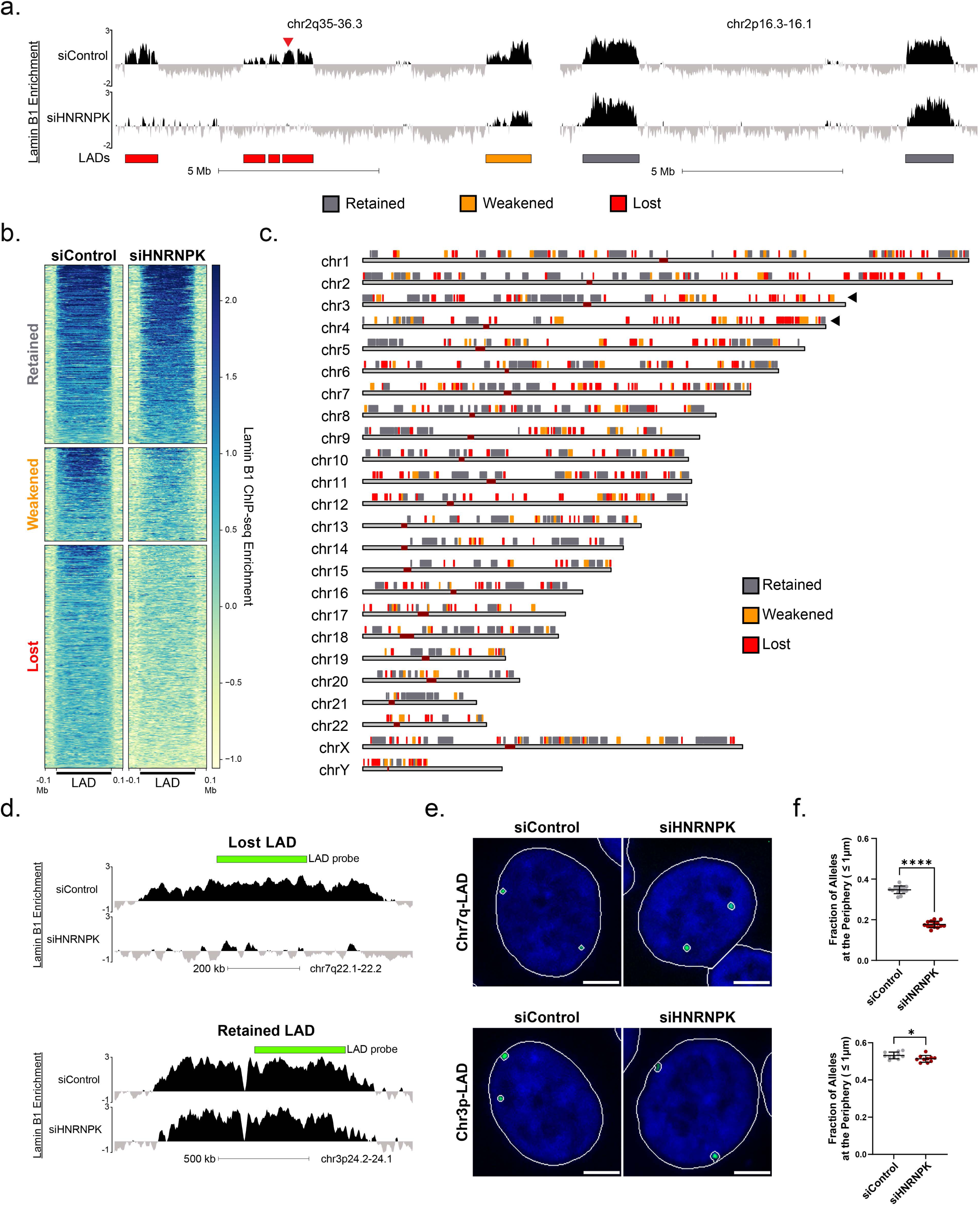
hnRNPK regulates LAD organization genome-wide. a) Lamin B1 ChIP-seq enrichment across chr2q35-36.3 (left) and chr2p16.3-16.1 (right) in control or *HNRNPK* KD. Red arrow indicates screened Chr2q-LAD. Bottom track represents control-defined LADs colored by sensitivity to hnRNPK loss-retained (gray), weakened (orange), lost (red). b) Heatmap of Lamin B1 enrichment [*z*-score log_2_(IP/input)] across control-defined LADs grouped by sensitivity to hnRNPK loss. Domains scaled to the same size with fixed 100 kb flanking regions. c) Ideogram showing chromosomal distribution of control-defined LADs colored by sensitivity to hnRNPK loss. Maroon bands represent centromeres. Black arrows indicate clusters of lost and weakened LADs on specific chromosome arms. d) Lamin B1 ChIP-seq enrichment in control and *HNRNPK* KD and Oligopaint probe design (indicated in green) at a lost (top) and retained (bottom) LAD on chr7q22.1-22.2 and chr3p24.2-24.1 respectively. e) Representative DNA HiDRO images of a lost (Top, Chr7q-LAD) and a retained (Bottom, Chr3p-LAD) LAD in control and *HNRNPK* KD nuclei. Solid white line indicates nuclear edge defined by DAPI staining (blue). Scale bar, 5 µm. f) Quantification of DNA HiDRO analyses showing fraction of alleles within 1 µm of the nuclear periphery for a lost (Top, Chr7q-LAD) and a retained (Bottom, Chr3p-LAD) LAD in control and *HNRNPK* KD. Each dot represents an individual HiDRO well. >300 nuclei analyzed per well. Error bars represent mean and SD of replicates. Chr7q-LAD (12 wells per condition), **** *P =* <0.0001; Chr3p-LAD Rep. 1 (10 wells per condition), * *P* = 0.036. Unpaired t test with Welch’s correction.

LADs showed a spectrum of sensitivity to hnRNPK depletion across the genome. While many LADs were lost, similar to our screened locus, others were weakened, and some remained unchanged. Notably, virtually no de novo LADs formed following hnRNPK depletion (Supp. Fig. 3b). To quantify these changes, we classified LADs identified in control cells into three categories based on their fractional base pair overlap with LADs in the siHNRNPK condition: retained (75–100% overlap), weakened (25–75% overlap), and lost (0–25% overlap, Fig. 3a-b, Supp. Fig. 3c-d, Supplementary Table 5). Using these criteria, we identified 317 retained LADs, 166 weakened LADs, and 396 lost LADs, indicating that ∼64% of LADs, spanning 309 Mb, are sensitive to hnRNPK depletion (Supp. Fig. 3e). In addition, approximately 30% of retained LADs, which were significantly larger than weakened and lost LADs, displayed selective loss of Lamin B1 at their edges (Supp. Fig. 3f-g), suggesting border-specific modulation of lamina association in a subset of these domains as well.

Finally, hnRNPK-sensitive LADs were distributed across all chromosomes (Fig. 3c), though we observed regional clustering in some cases; for example, the q arms of chromosomes 3 and 4 harbored a dense cluster of weakened and lost LADs, whereas chromosome 3p and 4p displayed relatively few hnRNPK-sensitive domains (Fig. 3c, black arrows). To validate these site-specific effects, we used Oligopaint DNA FISH to examine the spatial positioning of selected LADs in control and siHNRNPK cells (Fig. 3d). Indeed, a lost LAD on chromosome 7q showed a ∼50% decrease in perinuclear localization frequency upon hnRNPK knockdown (Fig. 3e-f, Supp. Fig. 3h-i). In contrast, a retained LAD on chromosome 3p remained peripherally positioned, with minimal change in perinuclear distance, supporting the specificity of the ChIP-seq results (Fig. 3e-f, Supp. Fig. 3h-i).

### hnRNPK sensitivity defines two epigenetically and functionally distinct classes of LADs

To investigate what distinguishes hnRNPK-sensitive LADs from domains that are unaffected by its loss, we profiled key heterochromatin-associated histone post-translational modifications by ChIP-seq. Highly concordant replicates (n ≥ 2 per condition) were merged for analysis (Supp. Fig. 4a). We first examined enrichment of these modifications in the control condition, focusing on LADs stratified by their sensitivity to hnRNPK knockdown. As expected, the perinuclear marker H3K9me2 was enriched across nearly all LADs, though retained LADs demonstrated greater H3K9me2 enrichment compared to weakened or lost LADs in control cells (Supp. Fig. 4b).

In contrast, the distribution of H3K9me3 and H3K27me3 revealed striking differences between hnRNPK-sensitive and -insensitive LADs (Fig. 4a-b). LADs retained following hnRNPK depletion were enriched for H3K9me3 across their domain bodies, and H3K27me3 enrichment was restricted to their borders. Conversely, LADs that were lost or weakened upon hnRNPK KD were depleted of H3K9me3 and instead enriched for H3K27me3 across the entire domain (Fig. 4b). This mutually exclusive enrichment pattern accounted for nearly all LADs genome-wide, with 479 Mb co-occupied by Lamin B1 and H3K9me3 and 314 Mb co-occupied by Lamin B1 and H3K27me3 (Fig. 4c). It is important to note, however, that a substantial fraction (420 Mb) of H3K27me3-enriched chromatin is found outside of LADs (Fig. 4c), consistent with its broader roles in genome regulation^60^.

**Figure 4.**
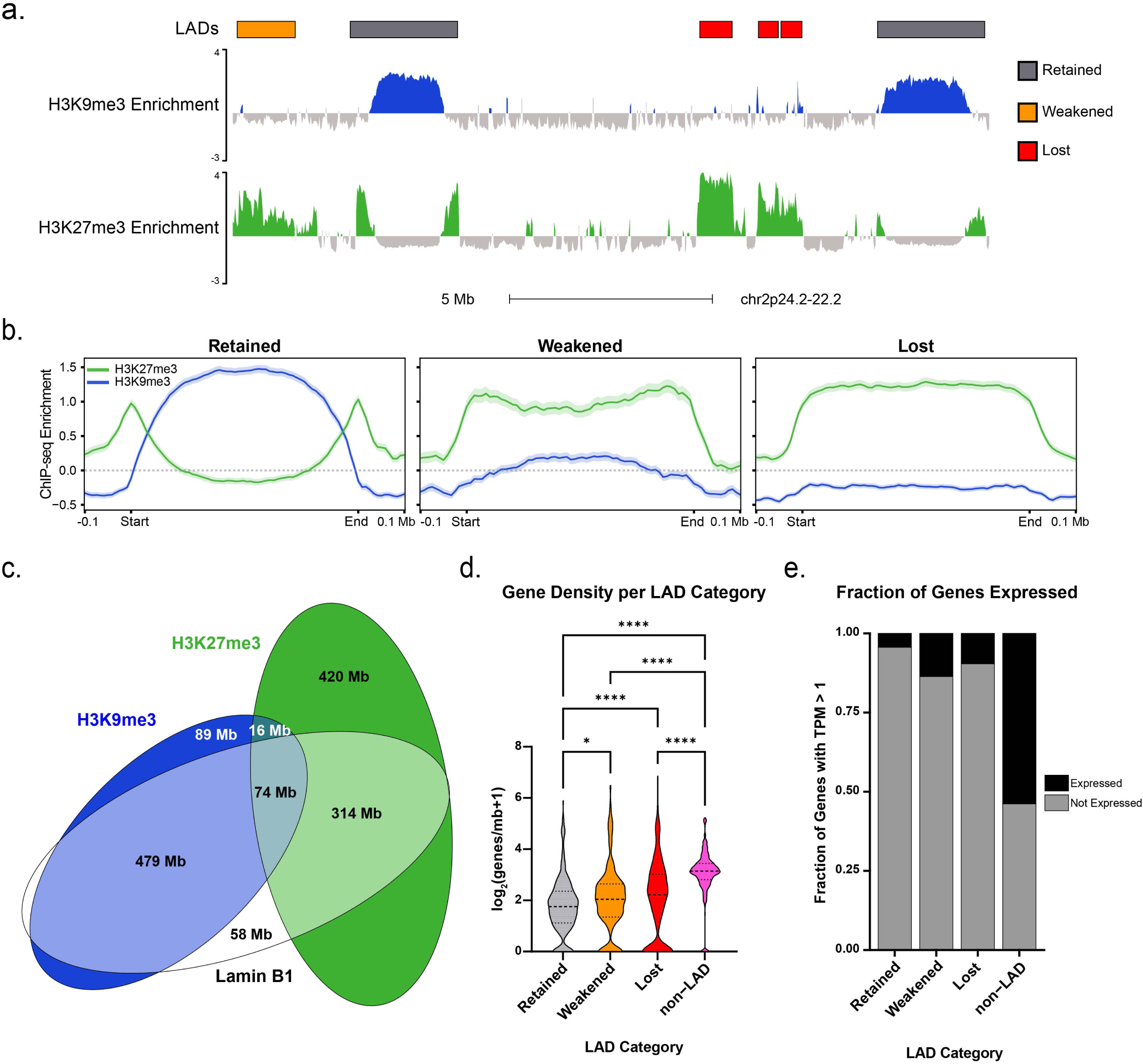
hnRNPK sensitivity defines two epigenetically and functionally distinct classes of LADs. a) H3K9me3 (blue) and H3K27me3 (green) ChIP-seq tracks of chr2p24.2-22.2 in RNAi siControl. Top track represents control-defined LADs colored by sensitivity to hnRNPK loss. b) H3K9me3 (blue) and H3K27me3 (green) occupancy [*z*-score log_2_(IP/input)] in siControl at retained, weakened and lost LADs. Domains scaled to the same size. c) Venn diagram of base pair overlap between H3K9me3 (blue), H3K27me3 (green) and Lamin B1 (white) domains. d) Gene density in retained (median 2.4 genes/mb), weakened (median 3.1 genes/Mb), lost LADs (median 3.6 genes/Mb) and nonLAD regions (median 7.8 genes/Mb). Boxplot represents 1^st^ quartile, median and 3^rd^ quartile. Kruskal Wallis followed by Dunn’s test. Retained vs. weakened (*P* = 0.03); retained vs. lost (p.adj ≤ 0.0001); weakened vs. lost ns (*P* = 0.3); all LAD categories vs. non-LAD (*P* ≤ 0.0001). e) Fraction of genes expressed (TPM >1) in retained LADs (4.4% expressed), weakened LADs (13.7% expressed), lost LADs (9.6% expressed) and nonLADs (53.8% expressed).

To further examine whether hnRNPK-sensitive LADs differ from insensitive LADs, we examined their gene content and transcriptional activity in control cells by RNA-seq. Although lost and weakened LADs are significantly more gene-dense than retained LADs on average, both classes remain gene-poor relative to nonLAD regions (Fig. 4d). Importantly, hnRNPK-sensitive LADs exhibit similarly low transcriptional activity as retained LADs. Only 4–14% of genes are expressed across all LAD categories in control cells, in stark contrast to ∼50% of genes expressed in nonLAD regions (Fig. 4e). Moreover, genes within LADs are expressed at significantly lower levels than genes outside LADs (Supp. Fig. 4c), indicating that hnRNPK-sensitive LADs are not inherently more transcriptionally active or “less LAD-like” than retained LADs. Gene ontology analysis revealed that lost LADs are enriched for developmental and differentiation-related genes, whereas retained LADs are enriched for housekeeping functions such as cell junction assembly and cell-cell adhesion (Supp. Fig. 4d). These findings support the existence of two mechanistically and epigenetically distinct classes of LADs, with hnRNPK as a selective regulator of the H3K27me3-enriched class of perinuclear heterochromatin.

### LAD repositioning upon hnRNPK loss derepresses a subset of genes

We next asked whether LAD detachment following hnRNPK depletion alters the heterochromatin or transcriptional state of these regions. Western blotting and immunofluorescence analysis of global histone post-translational modifications levels showed no significant change in levels or nuclear localization of H3K9me3 or H3K27me3 following 72-hour hnRNPK KD (Supp. Fig. 5a-b). We then analyzed the enrichment of these modifications specifically over hnRNPK-sensitive LADs. Despite their reduced lamina association, lost LADs remained enriched for H3K27me3 and depleted of H3K9me3 in siHNRNPK cells (Fig. 5a). Similar results were observed for weakened LADs (Supp. Fig. 5c).

**Figure 5.**
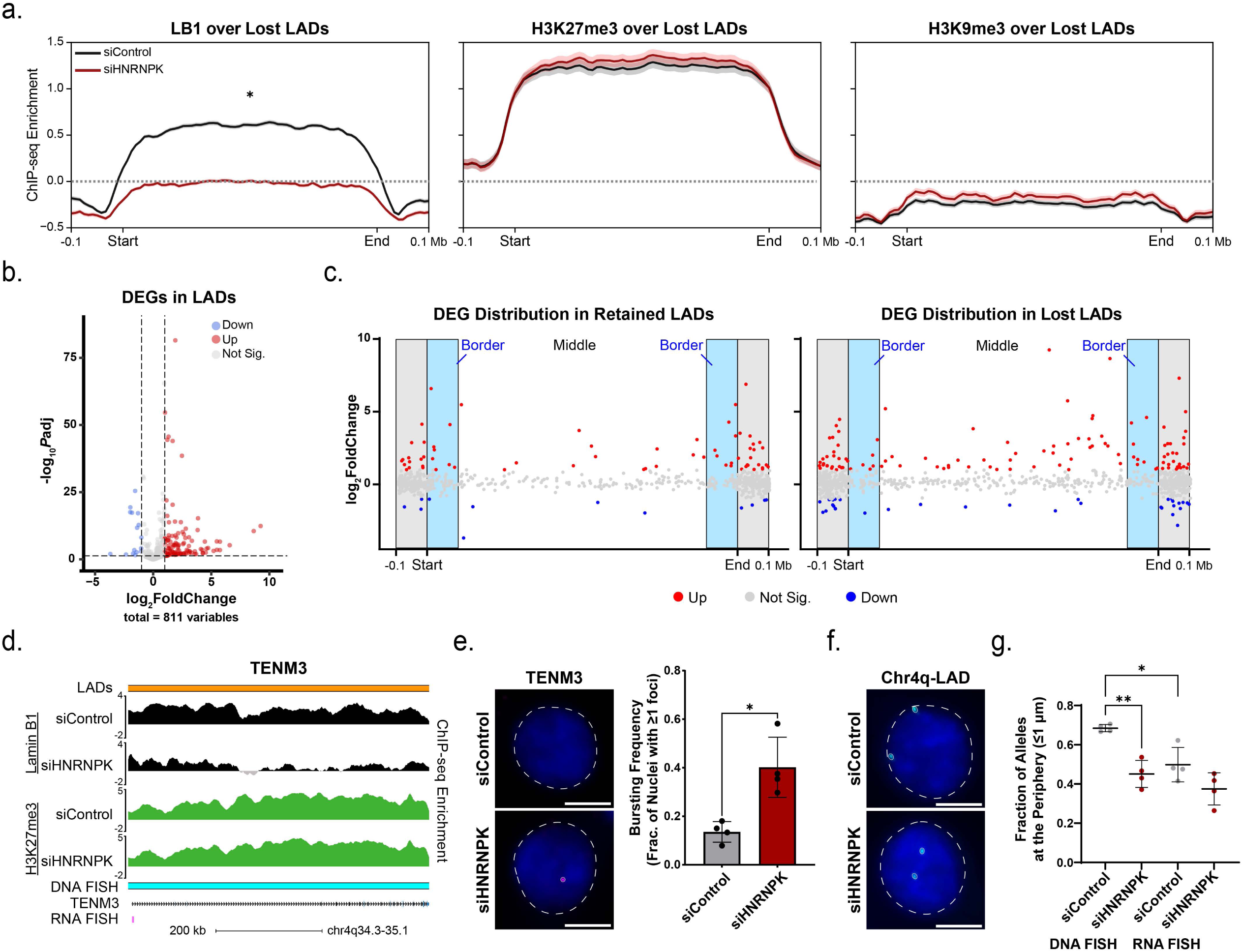
Transcriptional effects of LAD repositioning upon hnRNPK loss. a) Lamin B1 (left), H3K27me3 (middle), H3K9me3 (right) enrichment [*z*-score log_2_(IP/input)] over lost LADs in control (black) and si*HNRNPK* (dark red) KD. Domains are scaled to the same size. Statistical analysis was performed using Wilcoxon signed-rank tests with threshold *P* < 0.05: Lamin B1 siControl vs siHNRNPK (passed *P* < 0.05 threshold), H3K27me3 siControl vs siHNRNPK (ns), H3K9me3 siControl vs siHNRNPK (ns). b) Volcano plot of genes within siControl-defined LADs after *HNRNPK* KD versus significance, denoting upregulated DEGs (red, n = 123), downregulated DEGs (blue, n = 16), and non-DEGs (gray, n = 672). Dotted lines indicate thresholds (adj. *P* < 0.05; |log_2_FC| > 1) c) The relative position of genes within retained (left) and lost (right) LADs versus their log_2_ fold change after *HNRNPK* KD. DEG status is denoted by color with upregulated genes labeled red, downregulated labeled blue, and non-DEGs labeled black. Light blue shaded rectangle represents the border region within LADs. Grey shaded rectangle represents fixed 100 kb flanking regions. d) Representation of Oligopaint RNA-FISH probe (indicated in magenta) targeting *TENM3* in a weakened LAD (indicated in orange) on chromosome 4 (chr4q34.3-35.1). Lamin B1 (black) and H3K27me3 (green) ChIP-seq tracks for control and *HNRNPK* KD are shown. A portion of the Oligopaint DNA-FISH probe targeting the LAD containing *TENM3* is shown in cyan. e) [Left] Representative images of intronic RNA FISH to the *TENM3* transcript (red) in control and *HNRNPK* KD nuclei. Dashed white line indicates nuclear edge as defined by DAPI staining (blue). Scale bar, 5 μm. [Right] Quantification of 3D RNA FISH analyses showing the bursting frequency of *TENM3* in control and *HNRNPK* KD. Bursting frequency was determined by calculating the fraction of nuclei with ≥ 1 RNA FISH foci. Each dot represents one biological replicate. > 500 nuclei analyzed per replicate. * *P* = 0.018, Unpaired t test with Welch’s correction. f) Representative images of 3D DNA FISH to the Chr4q-LAD (green) that overlaps *TENM3* in control and *HNRNPK* KD nuclei. Dashed white line indicates nuclear edge defined by DAPI staining (blue). Scale bar, 5 μm. g) Quantification and comparison of 3D DNA and RNA FISH analyses showing the fraction of either Chr4q LAD (DNA, left) or intronic *TENM3* (RNA, right) alleles within 1 μm of the nuclear periphery in control (black) and *HNRNPK* KD (dark red). Each dot represents an individual biological replicate, > 300 nuclei analyzed per replicate. Error bars represent mean and SD of replicates. siControl-DNA vs. siHNRNPK-DNA, \*\**P =* 0.021; siControl-DNA vs. siControl-RNA, \**P =* 0.022; siControl-RNA vs. siHNRNPK-RNA, ns *P* = 0.086. Unpaired t test with Welch’s correction.

Comparing gene expression between siControl and siHNRNPK samples, we identified 2,448 differentially expressed genes (DEGs) from RNA-seq datasets, with a similar number of upregulated (1,324) and downregulated (1,124) genes (Supp. Fig. 5d). This is consistent with the many functions of hnRNPK in both transcriptional activation and repression^61–63^. Saliently, integrating our gene expression datasets with our Lamin B1 ChIP-seq revealed proportionally more upregulated genes within LADs as compared to nonLADs, with 16-17% of genes becoming upregulated in hnRNPK-sensitive domains (Fig. 5b, Supp. Fig. 5e). Moreover, retained LADs showed a significant depletion of DEGs in the middle of their domains, consistent with their border-specific loss of Lamin B1 (Fig. 5c, Supp. Fig. 5f-g). In contrast, DEGs found in weakened and lost LADs were distributed evenly across the domains (Fig. 5c, Supp. Fig. 5f-h). Notably, most hnRNPK-sensitive LADs did not contain any upregulated DEGs (Supp. Fig. 5j), and 15-29% of LADs lack an annotated TSS (Supp. Fig. 5i). Thus, it is unlikely that local changes in transcription following hnRNPK knockdown are sufficient to induce LAD repositioning.

To further examine the relationship between lamina association and transcriptional activity, we performed intronic RNA FISH for a representative gene, *TENM3*, which resides in a hnRNPK-sensitive LAD on chr4q and is identified as significantly upregulated in the RNA-seq dataset (Fig. 5d). No change in H3K27me3 levels was observed across the gene (Fig. 5d). However, quantification of intronic RNA foci revealed a significant increase in transcriptional bursting frequency (∼15% to ∼40%) following hnRNPK KD (Fig. 5e), with no detectable change in burst amplitude (RNA signal intensity per active allele, Supp. Fig. 5k). These results validate its aberrant activation and suggest this occurs through more frequent transitions into the active state rather than stronger bursts. Finally, we compared the spatial positioning of *TENM3* transcripts by intronic RNA FISH with that of DNA FISH for the same locus. In control cells, intronic RNA signal was located further from the nuclear periphery than their corresponding DNA loci (Fig. 5f-g), suggesting that transcriptional activation occurs preferentially when alleles are further from the lamina. Together, these results support a model in which hnRNPK loss leads to LAD detachment from the nuclear periphery, which in turn drives increased transcriptional activity in a subset of previously silenced genes, potentially through enhanced bursting frequency.

## Discussion

Using the first genome-wide DNA FISH screen for chromatin positioning, we uncovered a previously unappreciated requirement for hnRNPK in maintaining the peripheral localization of a specific subset of LADs in human cells (Fig. 6). LADs that are sensitive to hnRNPK depletion are strongly enriched for H3K27me3 and depleted of H3K9me3, whereas LADs that are retained following hnRNPK knockdown display the opposite pattern. This mutually exclusive pattern of enrichment supports the existence of at least two mechanistically and epigenetically distinct classes of LADs. These observations are consistent with prior reports describing variation in LAD chromatin features, including those that distinguish constitutive and facultative LADs^11–15^. Our results build on this model by identifying hnRNPK as a selective regulator of the H3K27me3-enriched LAD subset. In contrast, LADs enriched for H3K9me3 may be maintained through distinct mechanisms, such as PRR14-mediated tethering, which has been implicated in anchoring H3K9me3 heterochromatin to the lamina in both mouse and human cells^40,43,64^. Together, these findings support a model in which LAD organization is governed by multiple, possibly redundant, pathways that differentially engage chromatin subtypes to preserve nuclear architecture and gene repression.

**Figure 6:**
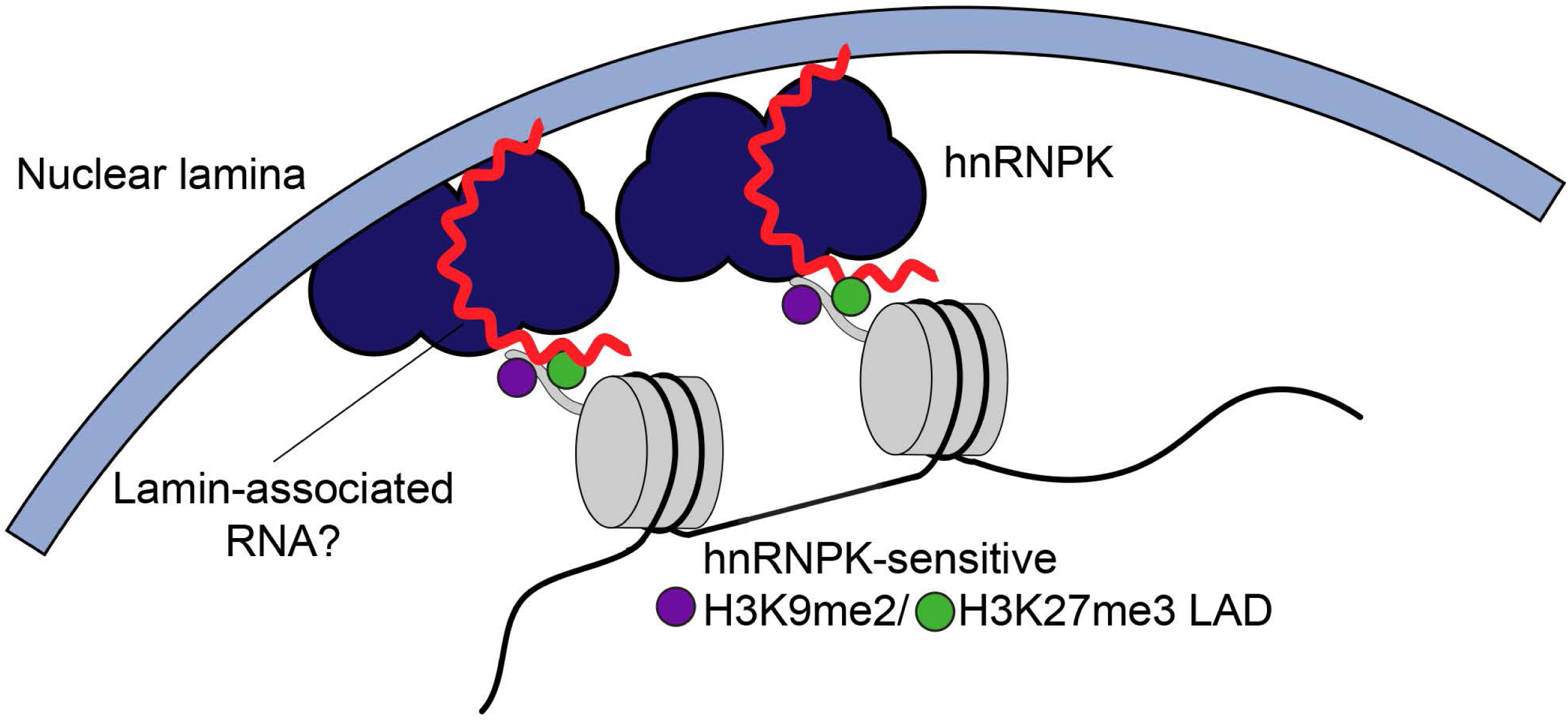
Model. Schematic illustrating hnRNPK as a putative peripheral tether for LADs marked by both H3K9me2 and H3K27me3. In this model, a lamina-associated RNA (red) interacts with hnRNPK (dark blue), recruiting it to the nuclear lamina and/or directly to LAD chromatin. This RNA-mediated recruitment is proposed to confer specificity to hnRNPK’s tethering function, thereby stabilizing the peripheral positioning and correct silencing of this LAD subclass.

While hnRNPK is known to function broadly in transcriptional regulation and RNA metabolism, several lines of evidence suggest that its effect on LAD positioning is unlikely to reflect general perturbations to these processes. Most notably, hnRNPK was the only member of the hnRNP family identified in our genome-wide screen, and we did not observe significant enrichment for general components of the transcriptional or splicing machinery. These findings argue for a selective role for hnRNPK in genome organization, distinct from its broader molecular functions. At the molecular level, our identification of hnRNPK suggests a model in which RNA participates in anchoring heterochromatin at the nuclear periphery (Fig. 6). In further support, PRMT1, an arginine methyltransferase known to regulate hnRNPK’s RNA-binding activity^65–68^, also emerged as a hit in our screen. These findings raise the possibility that specific RNA species, transcribed from LADs or acting in *trans*, recruit hnRNPK to its genomic targets, providing specificity to this unique function and analogous to how the lncRNA Xist recruits hnRNPK to the inactive X chromosome^57,69^. Indeed, the inactive X is similarly marked by H3K27me3 and localized to the nuclear periphery^70^. Given the robust LAD repositioning observed upon hnRNPK depletion and the enrichment of additional RNA-processing proteins among our screen hits, we speculate that additional RNA-processing factors or RNA itself may function in parallel or in concert with hnRNPK to organize peripheral heterochromatin in human cells. This model is consistent with prior studies implicating RNA-binding proteins in both the heterochromatin and lamina-associated proteomes, as well as the role of low-level transcription in heterochromatin maintenance^13,56,71^. Whether these factors function independently or as multi-protein RNA–chromatin complexes remains to be determined.

Surprisingly, despite large-scale changes in chromatin positioning, we observed no clear alterations in histone modification patterns following 72-hour depletion of hnRNPK. H3K27me3, in particular, remained enriched across LADs that were repositioned away from the nuclear periphery, indicating that peripheral localization is not strictly required for maintaining this repressive mark. Nonetheless, expression of a subset of genes within these repositioned LADs was altered, suggesting that chromatin state and transcriptional activity can become uncoupled from subnuclear localization. This finding is consistent with recent reports showing similar decoupling following depletion of multiple lamina components^26^. Although repositioned LADs were more likely to contain upregulated genes, the majority of genes within these domains remained transcriptionally silent. These results argue against transcriptional activation as the driver of LAD repositioning and instead support a model in which hnRNPK-mediated anchoring acts upstream of transcriptional regulation. This aligns with prior studies showing that movement away from the nuclear periphery is permissive but not always sufficient to activate transcription^7,8,21,22,24,72,73^.

Taken together, our work establishes HiDRO as a powerful platform for spatial genome-wide screening and a conceptual framework for understanding chromatin-lamina interactions. We define a unique mechanism by which hnRNPK mediates facultative heterochromatin marked by H3K27me3 at the nuclear periphery. This study opens the door to future investigations of LAD subclasses, the role of RNA in peripheral chromatin organization, and the potential for chromatin-lamina architecture to be dynamically remodeled in development and disease.

---

**Supp. Figure 1.**
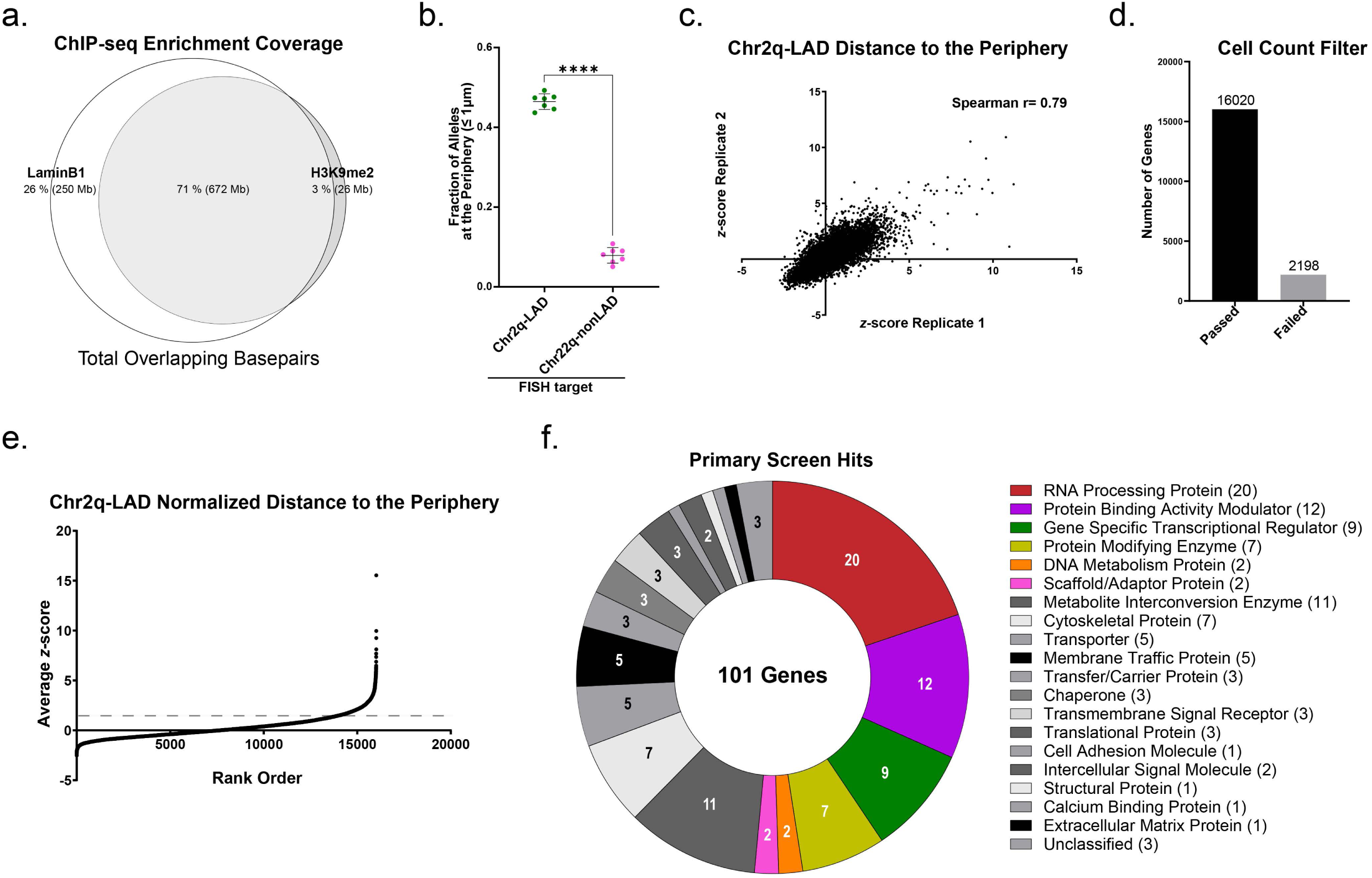
a) Venn diagram of base pair overlap between control LADs and H3K9me2-domains in HCT116 cells. b) Quantification of DNA HiDRO analyses (Fig. 1c) showing the fraction of alleles within 1 μm of the nuclear periphery for the Chr2q-LAD (green) and the Chr22q-nonLAD (magenta). Each dot represents an individual HiDRO well (>500 alleles per well). Error bars represent mean and SD of replicate wells. **** *P* <0.0001, Unpaired t test with Welch’s correction. c) Correlation of robust *z*-scores for the Chr2q-LAD Euclidean distance to the periphery between biological replicates of the primary screen. Only genes that passed cell count threshold are shown. Spearman’s correlation r= 0.79. d) Bar chart showing the number of genes that passed (black, n = 16,020) and failed to meet (gray, n = 2,198) the cell count threshold in the primary HiDRO screen. e) Rank-ordered robust *z*-scores for the normalized distance to the periphery for the Chr2q-LAD for all genes passing the cell count threshold. *Z*-scores are the average of two biological replicates. f) Protein classes of primary hits of HiDRO screen. Colored groups indicate classes tested in validation screen. n = 101 genes.

**Supp. Figure 2.**
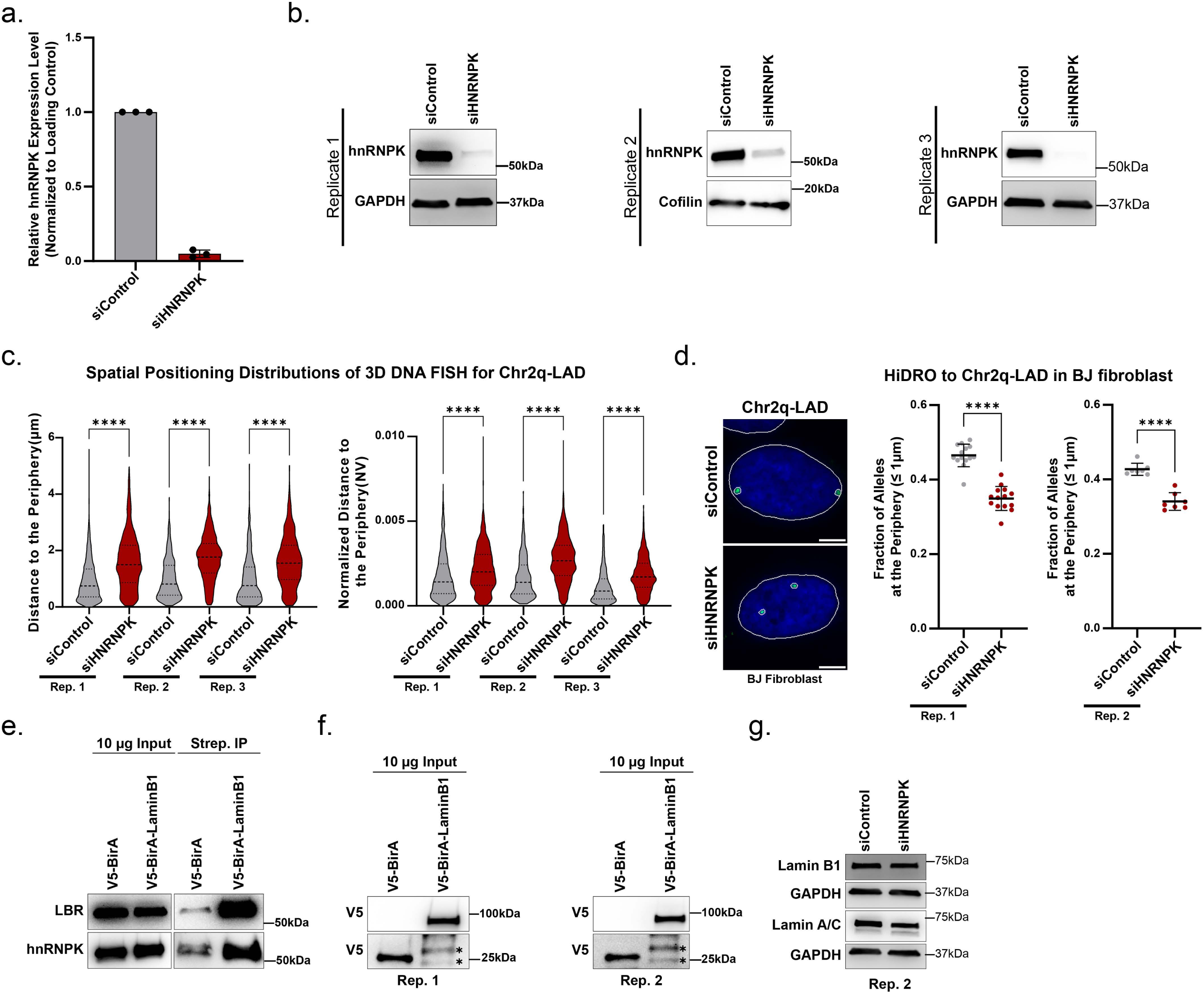
a) Quantification of hnRNPK levels relative to control following *HNRNPK* KD (blots shown in Supp. 2b). Data are the mean of three biological replicates. Error bars represent mean and SD of replicates. b) Western blot showing global levels of hnRNPK in from control and *HNRNPK* KD whole cell lysate from three biological replicates. GAPDH and Cofilin used as loading controls. c) (Left) Quantification of 3D DNA FISH analyses showing the distributions of distance to the nuclear periphery of the Chr2q-LAD in control (gray) and *HNRNPK* KD (dark red) across 3 biological replicates (shown in Fig. 2C). (Right) Quantification of 3D DNA FISH analyses showing the distributions of normalized distance to the nuclear periphery of the Chr2q-LAD in control (gray) and *HNRNPK* KD (dark red). > 600 alleles analyzed per replicate. Dotted lines in violin plots denote 1^st^ and 3^rd^ quartile, dashed line represents the median. **** *P* <0.0001, Kruskal-Wallis test with uncorrected Dunn’s test. d) (Left) Representative DNA HiDRO image in BJ fibroblasts of the Chr2q-LAD (green) in control and *HNRNPK* KD nuclei. Solid white line indicates nuclear edge defined by DAPI staining (blue). Scale bar, 5 μm. [Right] Quantification of DNA HiDRO analyses in BJ fibroblasts showing the fraction of Chr2q-LAD alleles within 1 μm of the nuclear periphery in control (black) and *HNRNPK* KD (dark red). Each dot represents an individual HiDRO well across 2 independent HiDRO plates (Right Rep. 1 and Left Rep.2). > 200 nuclei and > 300 alleles analyzed per well. Rep.1 (14 wells per condition), **** *P =* <0.0001; Rep. 2 (7 wells per condition), **** *P =* <0.0001. Unpaired t test with Welch’s correction. e) Additional replicate of western blot for Lamin B receptor and hnRNPK in HCT116 cells after Turbo-ID biotin labeling from input (left) and streptavidin pull-down (right) from chromatin fractions. Corresponds to Fig. 2d. f) Western blot for V5 expression in HCT116 cells from input chromatin fraction for 2 replicates (left, right). Corresponds to blots in Fig. 2d and Supp. Fig. 2e. Asterisks indicate non-specific bands at lower molecular weights in V5-BirA-LaminB1 lanes. g) Additional replicate of western blot showing levels of Lamin B1 and Lamin A/C in control and *HNRNPK* KD whole cell lysates. GAPDH was used as a loading control. Corresponds to Fig. 2e.

**Supp. Figure 3.**
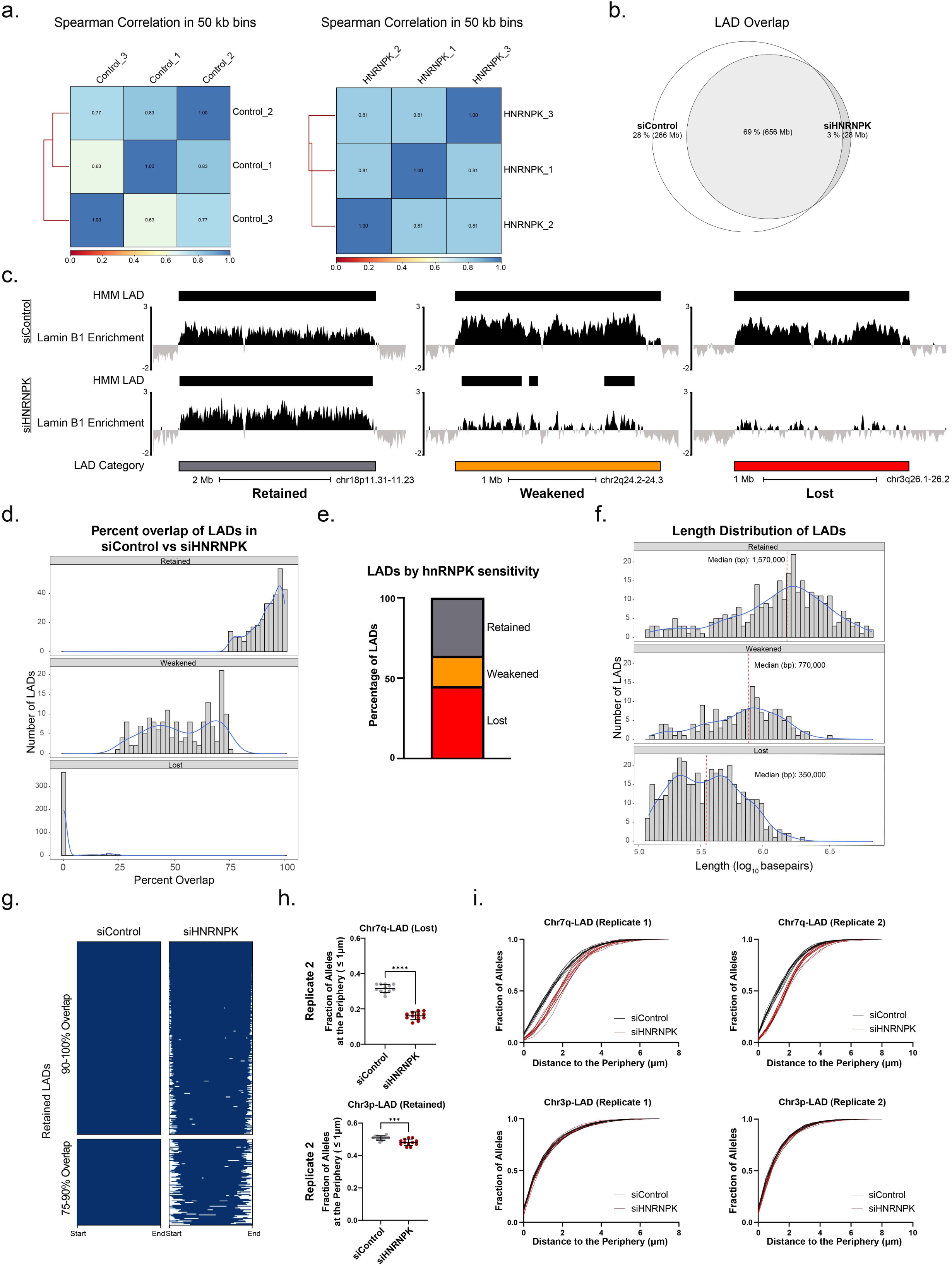
a) Spearman correlations among Lamin B1 ChIP-seq replicates (n=3) for siControl (left) and *HNRNPK* (right) KD, computed over 50kb windows. b) Venn diagram of total basepair overlap of LADs in siControl and *HNRNPK* KD. c) Lamin B1 ChIP-seq enrichment across representative retained, weakened or lost LADs in siControl (top) and *HNRNPK* (bottom) KD. Black bars above each track represent LAD calls in control or *HNRNPK* KD. Bars below tracks represent control LADs stratified by sensitivity to hnRNPK loss: retained (gray, 75-100% overlap between conditions), weakened (orange, 25-75%), lost (red, 0-25%). d) Distributions of percent overlap of LADs in siControl versus *HNRNPK* KD for retained (75-100% overlap), weakened (25-75%), and lost LADs (0-25%). e) Stacked bar graph showing percentage of siControl-defined LADs that are retained (36.1%, n=317), weakened (18.9%, n=166), or lost (45%, n=396) following *HNRNPK* KD. f) Size distributions of siControl LADs stratified by sensitivity to hnRNPK loss. Red dashed line indicates median in each group. Statistical test using Welch’s t−test with BH adjustment. Retained vs. weakened; retained vs. lost; weakened vs. lost, all comparisons **** *Padj*< 0.0001. g) Binary heatmap of retained LADs (domains scaled to the same size) in control and *HNRNPK* KD grouped by retained LADs with 90-100% vs 75-90% overlap. LAD indicated in blue and non-LAD in white. h) Quantification of an additional replicate of DNA HiDRO showing the fraction of alleles within 1 µm of the nuclear periphery for a lost (Top, Chr7q-LAD) and retained (Bottom, Chr3p-LAD) LAD in control and *HNRNPK* KD. Replicate corresponds to data in Fig. 3e,f. Each dot represents an individual HiDRO well. >300 nuclei analyzed per well. Error bars represent mean and SD of replicates. Chr7q-LAD Rep. 2 (14 wells per condition), **** *P =* <0.0001; Chr3p-LAD Rep. 2 (12 wells per condition), *** *P* = 0.0009. Unpaired t test with Welch’s correction. i) Cumulative distributions of distance to the nuclear periphery for a lost (Top, Chr7q-LAD) and a retained (Bottom, Chr3p-LAD) LAD in control and *HNRNPK* KD. Distributions correspond to the HiDRO experiments in Fig. 3e,f and Supp. Fig. 3h. Each curve represents an individual well, >300 nuclei analyzed per well. Chr7q-LAD Rep. 1 (Top left); Chr7q-LAD Rep. 2 (Top right); Chr3p-LAD Rep. 1 (Bottom left); Chr3p-LAD Rep. 2 (Bottom right). At least 10 wells per condition were analyzed.

**Supp. Figure 4.**
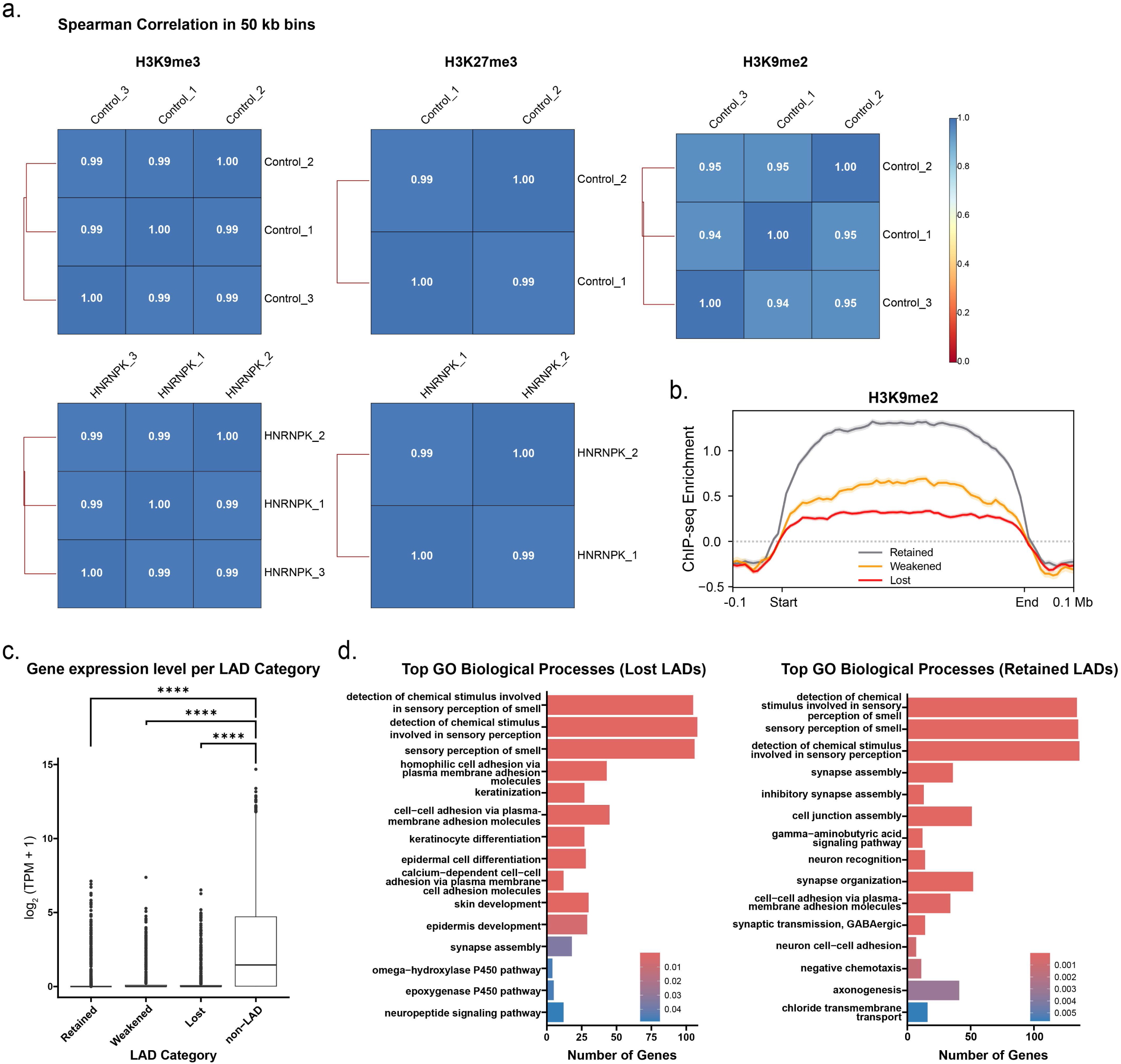
a) Spearman correlations across replicates for H3K9me3 (left), H3K27me3 (center), and H3K9me2 (right) ChIP-seq datasets in control and *HNRNPK* KD, computed over 50kb windows. b) H3K9me2 occupancy [*z*-score log_2_(IP/input)] at retained (gray), weakened (orange), and lost LADs (red) in siControl cells. Domains scaled to the same size. c) Comparison of control gene expression levels between retained LADs (median 0 TPM), weakened LADs (median 0 TPM), lost LADs (median 0 TPM), and non-LADs (median 1.74 TPM). Boxplot represents 1^st^ quartile, median and 3^rd^ quartile. All LAD categories had significantly lower gene expression than non-LADs, p.adj<0.0001 Kruskal-Wallis followed by Dunn’s test for multiple comparisons. d) Gene ontology enrichment analysis (biological processes) for genes within lost LADs (left) and retained LADs (right). Bars colored by significance (adjusted p-value).

**Supp. Figure 5.**
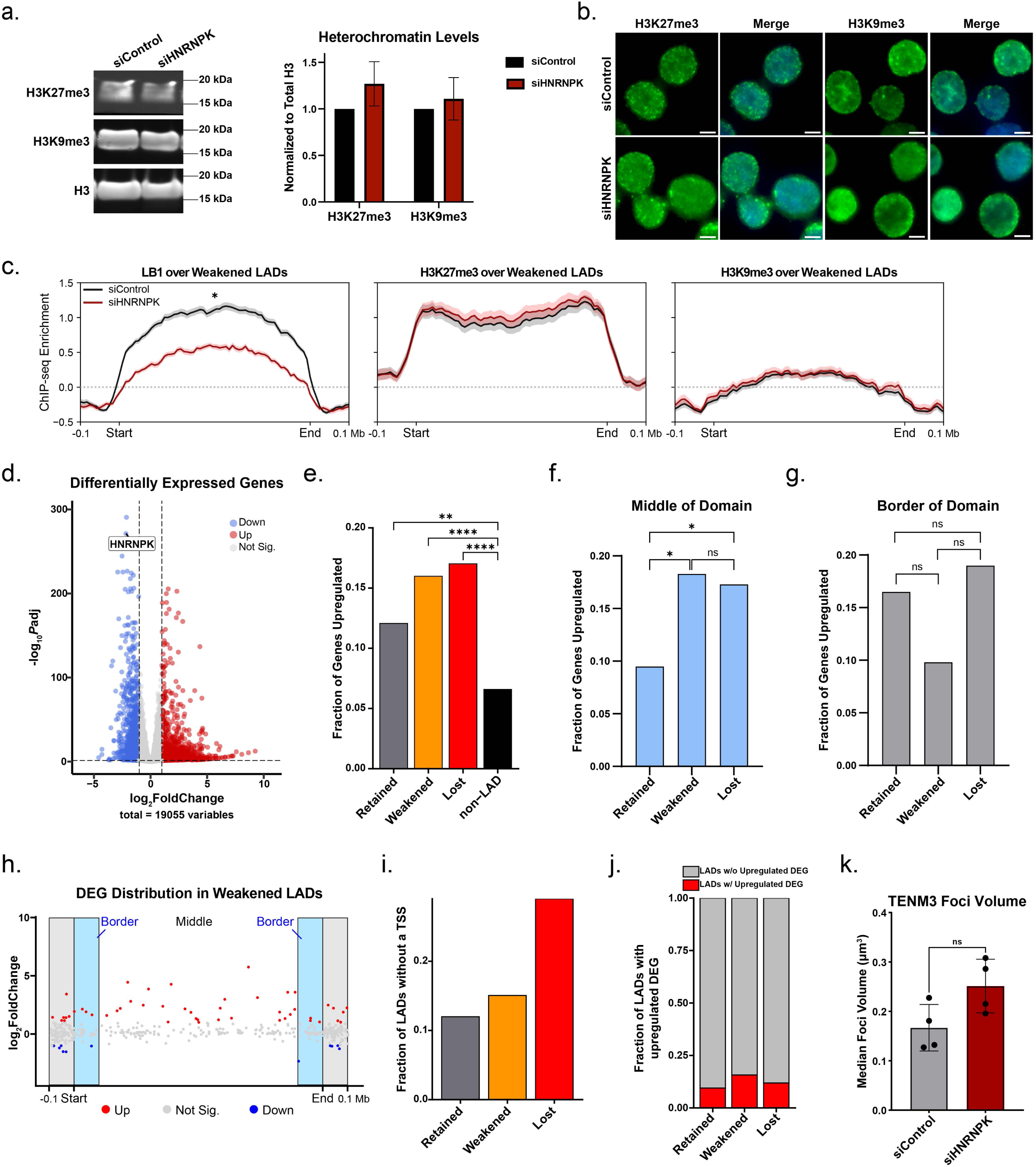
a) (Left) Western blot for total levels of H3K27me3 and H3K9me3 from chromatin fractionation of control and *HNRNPK* KD lysates. Total H3 is a loading control. (Right) Quantification of H3K27me3 and H3K9me3 levels relative to control following *HNRNPK* KD. Error bars represent mean and SD of replicates. H3K27me3 siControl vs siHNRNPK (ns *P* = 0.19), H3K9me3 siControl vs siHNRNPK (ns *P* = 0.50), unpaired t test with Welch’s correction. b) Immunofluorescence for H3K27me3 (green) and H3K9me3 (green) in siControl and *HNRNPK* KD nuclei. DNA is stained with DAPI (blue). Scale bars, 5 μm. c) Lamin B1 (left), H3K27me3 (middle), H3K9me3 (right) enrichment [*z*-score log_2_(IP/input)] across weakened LADs in siControl (black) and *HNRNPK* (dark red) KD. Domains scaled to the same size. Metaprofiles were compared using dsCompareCurves with paired Wilcoxon signed-rank tests (two-sided, *P* < 0.05 threshold) and 1,000 bootstrap resamples to estimate 95% CIs on paired differences: Lamin B1 siControl vs siHNRNPK (passed), H3K27me3 siControl vs siHNRNPK (ns), H3K9me3 siControl vs siHNRNPK (ns). d) The log_2_ fold change of all genes after *HNRNPK* KD versus their significance. DEG status is denoted by color with upregulated (n = 1,324 genes) genes labeled red, downregulated (n=1,124 genes) labeled blue, and non-DEGs labeled gray (n = 16,607 genes). The dotted lines indicate thresholds used to define DEGs (adj. P < 0.05, |log_2_(FC)| > 1). e) Fraction of genes that are an upregulated DEG across retained (gray), weakened (orange), lost (red) LADs and nonLADs. Statistical analysis was performed using two-sided Fisher’s exact test, ** *P* < 0.01, **** *P* < 0.0001; retained-nonLAD *P* = 0.0014; weakened-nonLAD *P* < 0.0001; lost-nonLAD *P* < 0.0001. f) Fraction of genes that are upregulated and located within the middle (inner 80%) of retained, weakened, and lost LADs. Statistical analysis comparing enrichment of upregulated DEGs (n=14 for retained, n=28 for weakened, and n=42 for lost LADs) vs nonDEGs (n=133 for retained, n=125 for weakened, n=201 for lost LADs) was performed using two-sided Fisher’s exact test, * *P* < 0.05; retained-weakened \**P* = 0.03; retained-lost \**P* = 0.037; weakened-lost ns *P* = 0.79. g) Fraction of genes that are upregulated and located within the border (10% on each side for 20% total) of retained, weakened, and lost LADs. Statistical analysis comparing enrichment of upregulated DEGs (n=17 for retained, n=6 for weakened, and n=15 for lost LADs) vs nonDEGs (n=86 for retained, n=55 for weakened, n=64 for lost LADs) was performed using two-sided Fisher’s exact test; retained-weakened ns *P* = 0.26; retained-lost *P* = 0.70; weakened-lost ns *P* = 0.16. h) The relative position of genes within weakened LADs versus their fold change after *HNRNPK* KD. DEG status is denoted by color with upregulated genes labeled red, downregulated labeled blue, and non-DEGs labeled black. Light blue shaded rectangle represents the border region within LADs. Grey shaded rectangle represents a fixed 100 kb overhang region outside of LADs. i) Fraction of retained (gray) weakened (orange) and lost (red) LADs without an annotated TSS. j) Stacked bar chart showing the fraction of retained, weakened, and lost LADs that overlap an upregulated DEG (red bar) following *HNRNPK* KD. k) Quantification of 3D RNA FISH analyses showing the median foci volume of intronic RNA FISH to the *TENM3* transcript in control and *HNRNPK* KD. Each dot represents one biological replicate. > 500 nuclei analyzed per replicate. ns *P* = 0.06, Unpaired t test with Welch’s correction.

## Supplementary Table Legends

**Supplementary Table 1 Coordinates of domains labeled by Oligopaint probes.** List of the genomic coordinates in hg38 of all domains used for slideFISH (DNA and RNA) or DNA HiDRO in this study.

**Supplementary Table 2 Primary HiDRO screen data, related to** **Figure 1**. Table contains data for all genes tested in the primary HiDRO screen, including robust *z*-scores for the Euclidean and normalized distance to the periphery of the Chr2q-LAD and Chr22q-nonLAD probes, cell counts and robust *z*-scores for nuclear area. “Primary_screen_hit” is denoted as “TRUE” if a gene was considered a hit (see Methods).

**Supplementary Table 3 Primary HiDRO screen data for 101 hits, related to** **Figure 1**. Table contains data for the hits of the primary screen, including robust *z*-scores for the Euclidean and normalized distance to the periphery of the Chr2q-LAD and Chr22q-nonLAD probes, cell counts, and and robust *z*-scores for nuclear area also provided. Protein class and HiDRO validation screen summary data provided.

**Supplementary Table 4 Validation HiDRO screen data, related to** **Figure 2**. List of siRNA duplexes tested in validation HiDRO screen with data for the Euclidean and normalized distance to the periphery of the Chr2q-LAD. Cell counts and robust *z*-scores for nuclear area also provided. “Validated_Chr2q-LAD_Repositioning” is denoted as “1” if this duplex significantly increased the Chr2q-LAD’s Euclidean and normalized distance to the periphery in at least 2/3 replicates (see Methods).

**Supplementary Table 5 Control and siHNRNPK LADs.** LADs identified by HMM in siControl that were included in the hnRNPK sensitivity category designations. Size in siControl, size in siHNRNPK, and degree of overlap noted.

**Supplementary Table 6 Available siRNA oligonucleotide sequences used in this study.** This table contains the available sequences of siRNA used in this study for RNAi. For *HNRNPK,* target sequences administered as a pool.

**Supplementary Table 7 ChIP-seq read table**. This table contains sequencing statistics for all ChIP-seq experiments. Total number of uniquely mapping reads per condition (Lamin B1, H3K9me2, H3K9me3, H3K27me3 or input) per replicate.

## Acknowledgements

We thank all members of the Joyce and Jain laboratories for their comments at all steps of this work. We thank Harvard ICCB-Longwood Screening Facility for RNAi libraries, bioinformatics tools and other support for the screen. The work was funded by the National Institutes of Health (R35GM128903, EFJ; R01AG082437, EFJ, RJ; R35HL166663, RJ) and Burroughs Welcome Foundation (RJ).

## Materials and Methods

### Experimental Methods

#### Cell Culture

The HCT116 cell line was purchased from ATCC (RRID:CVCL_0291) and was cultured in McCoy’s 5A medium supplemented with 10% fetal bovine serum, 2 mM l-glutamine, 100 U ml-1 penicillin, and 100 µg ml-1 streptomycin at 37°C with 5% CO2.

The BJ fibroblast cell line was obtained from ATCC (RRID:CVCL_3653). Cells were cultured in Eagle’s Minimum Essential Medium supplemented with 10% fetal bovine serum, 100 U ml-1 penicillin, and 100 µg ml-1 streptomycin at 37°C with 5% CO2.

#### RNAi

RNAiMAX transfection reagent (Thermo Fisher Scientific) was diluted in Opti-MEM reduced serum medium (Thermo Fisher Scientific), and pipetted onto siRNA. The plates were incubated at room temperature for 20 min. HCT-116 cells were trypsinized and resuspended in antibiotic-free medium, then plated onto siRNA duplexes to obtain a final siRNA concentration of 25 nM. BJ fibroblasts were plated in a final siRNA concentration of 40 nM. After 72 h of RNAi treatment, cells were collected for experiments. Available siRNA sequences are provided in Supplementary Table 6.

#### Optimized Oligopaint design and synthesis

Oligopaints were designed to have 80 bases of homology using the OligoMiner ^74^ design pipeline with an average of 4 probes per kb and were purchased from Twist Bioscience. Probe coordinates are provided in Supplementary Table 1. Oligopaints were synthesized as described previously ^47^. Specifically, we directly labelled probes by dye conjugation. 60 nmol dye aliquots were made from either Alexa 488 (Thermo Fisher Scientific, A20000) or Alexa 647 (Thermo Fisher Scientific, A20006), which were resuspended in DMSO, then vacuum-desiccated and stored at −20 °C. 2nmol of probe was resuspended in 10 µl 0.3 M sodium bicarbonate, then mixed with 60 nmol of fluorescent dye. Probe was incubated at room temperature in the dark for at least 6 h to conjugate to dye. The probes were then purified using the Zymo DNA Clean & Concentrator-100 kit. A total of 90 µl water was added to the probe, then 200 µl of oligo binding buffer was added followed by 400 µl of ethanol. The probe mixture was vortexed and then transferred to a spin column, which was centrifuged at 11,000g for 30 s. The filter column was transferred to a new 1.5 ml tube and 100 µl water was added to elute the bound probe. After 1 min incubation at room temperature, the column was centrifuged at 16,000g for 1 min. The probe mixture was mixed again with 200 µl of oligo-binding buffer and 400 µl of ethanol and transferred to the same column as before, centrifuging at 11,000g for 30 s. The sample was washed twice with 600 µl wash buffer, centrifuging at 11,000g for 30 s and discarding the flowthrough each time. The column was spun-dry at 16,000g for 1 min, then transferred to a new 1.5 ml tube. Then, 100 µl of water was added to the column, incubated at room temperature for 1 min, then centrifuged at 16,000g for 1 min to elute the probe. The probe and dye concentration was determined using the Nanodrop spectrophotometer to confirm proper dye conjugation.

#### HiDRO for 384-well DNA FISH

HiDRO was performed as described previously ^47^. For experiments involving RNAi, 384-well plates (Perkin Elmer, 6057300) were either manually seeded with siRNA or ordered pre-seeded with siRNA (Dharmacon) from the Harvard Medical School ICCB-L. The following additional control siRNA (Dharmacon) was used: non-targeting control 5 (Supplementary Table 6). RNAiMAX transfection reagent (Thermo Fisher Scientific) was diluted in Opti-MEM reduced serum medium (Thermo Fisher Scientific) and pipetted onto siRNA using a manual multichannel pipette. The plates were then centrifuged and incubated at room temperature for 20 min. All centrifugations for HiDRO plates were performed at 1,200 rpm for 2 min at room temperature unless otherwise indicated. For most of the steps, pipetting was performed using the Matrix WellMate (Thermo Fisher Scientific). Cells were trypsinized and resuspended in antibiotic-free medium, then 1.5×10^3^ to 2.5×10^3^ cells were seeded in each well. Plates were centrifuged and incubated until collection for downstream experiments. On the day of collection, the medium was aspirated, PBS was added to all of the wells and the plates were centrifuged. PBS was aspirated and cells were fixed in each well with 4% PFA, 0.1% Tween-20 in 1× PBS for 10 min at room temperature. Plates were centrifuged once during fixation. The plates were then rinsed with 1× PBS and washed twice for 5 min with 1× PBS with a spin during each wash. Then, 70% ethanol was then added to each well, the plates were sealed with foil plate covers (Corning) and stored for at least 20 h at 4 °C before performing FISH. On the first day of DNA FISH, ethanol was aspirated and the plates were washed in 1× PBS for 10 min to reach room temperature. The plates were then centrifuged, washed briefly again in 1× PBS and centrifuged again. Cells were permeabilized for 15 min in 0.5% Triton X-100 and 5 min in 2× SSCT (0.3 M NaCl, 0.03 M sodium citrate and 0.1% Tween-20). Then 2× SSCT/50% formamide was added to all of the wells, and the plates were double sealed with foil covers. Predenaturation was performed at 85 °C for 3 min and then 60 °C for 20 min on heat blocks (VWR). The plates were then centrifuged, foil covers were removed and hybridization mix was added to the wells. The hybridization mix consisted of 50% formamide, 10% dextran sulfate, 4% polyvinylsulfonic acid (PVSA), 0.1% Tween-20, 2× SSC and each probe at 0.1 pmol µl −1. A total of 2 pmol of probe was used per 20 µl of hybridization mix. After centrifugation, the plates were double sealed with foil covers and denatured at 85 °C for 20 min on heat blocks. The heat blocks were covered to block light and preserve primary fluorescently labelled probes. The plates were centrifuged after denaturation and then hybridized overnight (20–24 h) at 37 °C. The next day, hybridization mix was aspirated and the plates were washed quickly twice with room temperature 2× SSCT, then with 60 °C 2× SSCT for 5 min. The plates were then washed with room temperature 2× SSCT for 5 min. Nuclei were stained by washing for 5 min in Hoescht (1:10,000 in 2× SSCT). The plates were centrifuged, washed for 15 min with 2× SSC and centrifuged again. The plates were mounted with 50 μl of imaging buffer (2× SSC, 10% glucose, 10 mM Tris-HCl, 0.1 mg ml−1 catalase, 0.37 mg ml−1 glucose oxidase) and centrifuged. Finally, the plates were mounted with 30 µl of mineral oil (Fisher Chemical), centrifuged, and sealed with foil covers. Plates imaged within 5 days of FISH.

#### 3D DNA slide FISH

FISH was performed as previously described ^75^ with minor adjustments. Cells were trypsinized and resuspended in fresh culture medium at 1 × 10^6^ cells per ml, then settled onto fibronectin-coated (Sigma Aldrich) glass slides for 2 h. Cells were fixed to the slides for 10 min with 4% paraformaldehyde in phosphate-buffered saline (PBS) with 0.1% Tween-20, followed by three washes in PBS for 5 min each wash. The slides were stored in PBS at 4 °C until use. On the first day of FISH, the slides were warmed to room temperature in PBS for 10 min then permeabilized in 0.5% Triton X-100 in PBS for 15 min with nutation. The slides were then incubated for 5 min each in 2× SSCT (0.3 M NaCl, 0.03 M sodium citrate and 0.1% Tween-20) and 2× SSCT/50% formamide at room temperature, followed by 1 h incubation in 2× SSCT/50% formamide at 37 °C. Hybridization buffer containing primary Oligopaint probes, hybridization buffer (10% dextran sulfate, 2× SSCT, 50% formamide and 4% polyvinylsulfonic acid (PVSA)), 5.6mM dNTPs and 10 µg RNase A was added to slides, covered with a coverslip, and sealed with rubber cement. Fifty pmol of probe was used per 25 µl hybridization buffer. Slides were then denatured on a heat block in a water bath set to 80 °C for 30min, then transferred to a humidified chamber and incubated overnight at 37 °C. The following day, the coverslips were removed and slides were washed in 2× SSCT at 60 °C for 15min and 2× SSCT at RT for 10min twice. Next, hybridization buffer (10% dextran sulfate, 2× SSCT, 10% formamide and 4% PVSA) containing secondary probes conjugated to fluorophores (10 pmol per 25 µl buffer) was added to slides, covered with a coverslip and sealed with rubber cement. Slides were placed in a humidified chamber and incubated for 2h at RT. Slides were washed in 2× SSCT at 60 °C for 15min and 2× SSCT at RT for 10min. To stain DNA, slides were washed with Hoechst (1:10,000 in 2× SSC) for 5min. Slides were washed in 2x SSC at RT for 5min. Slides were then mounted in SlowFade Gold Antifade (Invitrogen). For DNA slide FISH experiments involving primary labeled Oligopaint probes, 2pmol of probe was used per 25 µl hybridization buffer. After overnight incubation at 37 °C, coverslips were removed and slides were washed in 2× SSCT at 60 °C for 15min and twice in 2× SSCT at RT for 10min. To stain DNA, slides were washed with Hoechst (1:10,000 in 2× SSC) for 5min. Slides were washed in 2x SSC at RT for 5min. Slides were then mounted in SlowFade Gold Antifade (Invitrogen).

#### 3D RNA slide FISH

RNA FISH probes were designed as Oligopaint probes using oligo pools (OPools) from Integrated DNA Technologies. The transcripts were probed with RNA FISH Oligopaint probes designed with a similar OligoMiner pipeline used for DNA FISH, with the exception of using the default 36 to 41 nucleotide length range. Probes were ordered with an additional 32 bp attached to the 5’ end to enable detection of probes using secondary fluorescent oligos, ordered through IDT. Cells were seeded and fixed onto glass slides (Thermo Fisher Scientific), using the same fixation procedures as are used in DNA FISH. After fixation, cells were permeabilized overnight in 300µl of 70% ethanol containing 1% SDS, and RNA FISH was started within 1 week of cell fixation. On the first day of FISH, slides were washed once in 2× SSCT for 5 min, then hybridization mix containing 125 nM primary unlabelled Oligopaint probes, hybridization buffer (10% dextran sulfate, 2× SSCT, 50% formamide and 4% PVSA) and 1% SDS was added to slides, and the slides were covered with glass coverslips and sealed with rubber cement. The slides were then denatured for 3 min at 60 °C on a heat block in a water bath and incubated overnight at 37 °C in humidified chamber. The next day, the coverslips were removed and the slides were washed for 15 min each in prewarmed 60 °C 2× SSCT in a plastic coplin jar to avoid glass coplin jars cracking. The slides were washed twice for 5 min each in room temperature 2× SSCT, then secondary hybridization mix containing 50 nM secondary fluorescently labelled Oligopaint probes, hybridization buffer and 1% SDS was added to slides, and the slides were covered with glass coverslips and sealed with rubber cement. After 1 h incubation at 37 °C in a dark humidified chamber, the coverslips were removed and slides were then washed for 5 min 37 °C 2× SSCT. DNA was stained in Hoescht (1:10,000 in 37 °C 2× SSCT) for 5 min. The slides were then washed once for 3 min each in room temp. 2× SSCT, mounted with SlowFade Gold Antifade, and sealed under glass coverslips with nail polish. The slides were imaged within 24 h of completion of FISH.

#### Immunofluorescence

Cells were harvested via trypsinization, resuspended at 1e6 cells/mL in fresh medium and seeded on fibronectin-treated glass slides. Slides were spun at 1200 RPM for 5 minutes at room temperature and incubated at 37°C with 5% CO2 for 2 hours to allow cells to adhere. Slides were fixed in 4% paraformaldehyde for 10 minutes and then washed 3 times (5 minutes/wash) in PBS. Slides were stored at 4°C until use. For immunofluorescence, cells were permeabilized in 0.1% Triton-PBS for 15 minutes followed by 3 washes (10 minutes/wash) in PBST (PBS with 0.1% Tween-20). Cells were blocked in 1% BSA-PBST for 1 hour at room temperature. Primary antibodies were diluted in 1% BSA-PBST and then added to a glass coverslip that was adhered to the slide with rubber cement. The following concentrations of primary antibodies were used: H3K9me3 (Abcam ab8898, 1:500) and H3K27me3 (CST #9733, 1:1,000). Cells were incubated in primary antibody for 1 hour at room temperature. Coverslips were removed and cells were washed 3 times (10minutes/wash) in PBST. Secondary antibodies were diluted in 1% BSA-PBST and then added to a glass coverslip that was adhered to the slide with rubber cement.

The following concentrations of secondary antibodies were used: anti-Rabbit Alexa Fluor 647 (Jackson ImmunoResearch Labs 111-605-003, 1:1,000). Cells were incubated in secondary antibody for 1 hour at room temperature. Coverslips were removed, followed by 2 washes (10 minutes/wash) in PBST and 1 wash (5minutes/wash) in PBS. DNA was stained with Hoechst once (1:10,000 in PBS for 10 minutes) before transferring slides to PBS for mounting. Slides were mounted with SlowFade Gold Antifade Mountant (Invitrogen). Images were acquired on a Leica widefield fluorescence microscope, using a 1.4 NA ×63 oil-immersion objective (Leica) and Andor iXon Ultra emCCD camera.

For immunofluorescence in 384-well plates (Perkin Elmer, 6057300) the following protocol was followed. On the day of collection, the medium was aspirated, PBS was added to all of the wells and the plates were centrifuged. PBS was aspirated and cells were fixed with 4% PFA, 0.1% Tween-20 in 1× PBS for 10 min at room temperature. Plates were centrifuged once during fixation. Plates were washed 3x for 10 minutes with 1x PBS at room temp. with a spin during each wash. Cells were permeabilized for 5 minutes in 0.5% Triton-X-100 in 1x PBS. The plates were then washed twice for 5 minutes with 1x PBS with a spin during each wash. Cells were blocked in a blocking solution (1% BSA, 1% goat serum in PBST (0.1% Tween-20)) for 1hr at room temperature. The plates were sealed with foil covers during the blocking incubation period. Primary antibodies were diluted in 1% BSA in PBST (Lamin B1, Abcam ab16048, 1:1000; Lamin A/C, Santa Cruz sc376248 1:1,000) during blocking incubation. After blocking incubation, the plates were centrifuged, and block solution was aspirated. A manual multi-channel pipette was used to add 20 μl of primary antibody solution to all wells. Plates were sealed with foil covers and incubated for 1 hr at room temperature. Following incubation, primary antibody solution was aspirated and the plates were washed 3x for 5 minutes with PBST (0.1% Tween-20) with a spin during each wash. Secondary fluorescent antibodies were diluted in 1% BSA in PBST. The following concentrations of secondary antibodies were used: anti-Rabbit Cy5 (Abcam ab6564, 1:1,000), anti-Mouse Alexa Fluor Plus 555 (ThermoFisher A32727, 1:1,000). The final PBST wash was aspirated and 50 μl of secondary antibody solution was added to all wells with a manual multi-channel pipette. Plates were sealed with foil covers and incubated for 1 hr at room temperature. Secondary antibody solution was aspirated, and the plates were washed once for 10 min in PBST. The plates were washed 2 additional times with PBST for 5 mins. The plates were spun once during each spin. DNA was stained by washing the plate for 5 minutes in Hoechst (1:10,000 in 1x PBS). After aspirating Hoechst solution, the plates were washed once in 1x PBS for 5 min. Finally, the plate was mounted in imaging buffer (0.1x PBS (15mM NaCl), 2% glucose, 50 mM Tris-HCl, 0.1 mg ml−1 catalase, 0.37 mg ml−1 glucose oxidase) and centrifuged. Mineral oil was then added to each well, the plate was spun, and then sealed with two foil covers. Plates were imaged immediately following completion of immunofluorescence protocol.

#### Microscopy

Images for HiDRO (or immunofluorescence in 384-well plates) experiments were acquired on a Molecular Devices Image Xpress Micro 4 high-content microscope using a 0.95 NA ×60 air objective (Nikon) or a Molecular Devices Image Xpress Micro 4 Confocal high-content microscope with a 0.42 µm pinhole and 1.4 NA ×60 water-immersion objective. Max projections of z-series (6 images, 0.5 µm spacing) were generated automatically in MetaXpress and were used for downstream analyses.

Images of slides (DNA FISH, RNA FISH, and immunofluorescence) were acquired on a Leica widefield fluorescence microscope using a 1.40 NA ×63 oil-immersion objective with the Andor iXon Ultra emCCD camera.

#### ChIP and ChIP-seq Library Preparation

ChIP-seq was performed as previously described ^8,32^. HCT116 cells were crosslinked in culture by addition of methanol-free formaldehyde (Pierce, final 1% v/v) for 10 minutes at room temperature with gentle rotation. Crosslinking was quenched by addition of glycine (final 125 mM) for 5 minutes with gentle rotation. Media was discarded and replaced with PBS; cells were scraped and transferred to conical tubes and pelleted by centrifugation (1500 rpm, 3 minutes at room temperature). Resulting pellets were flash frozen in liquid nitrogen and stored at −80°C. For ChIP, 30μL protein G magnetic beads (per ChIP sample; ThermoFisher) were washed 3 times in blocking buffer (0.5% BSA in PBS); beads were resuspended in 250μL blocking buffer and 2μg antibody (Lamin B1, Abcam, ab16048; H3K9me2, Abcam, ab1220; H3K9me3, Abcam, ab8898; H3K27me3, Cell Signaling Technology, #9733) and rotated at 4°C for at least 6 hours. Crude nuclei were isolated from frozen crosslinked cells as follows: cell pellet (from 10cm plate or equivalent cell number) was resuspended in 10mL cold Lysis Buffer 1 (50mM HEPES-KOH pH7.5, 140mM NaCl, 1mM EDTA, 10% Glycerol, 0.5% NP-40, 0.25% Triton X-100, and protease inhibitors), and rotated at 4°C for 10 minutes, followed by centrifugation (1500 rpm at 4°C for 3 minutes). Supernatant was discarded and the pellet was resuspended in 10mL cold Lysis Buffer 2 (10mM Tris-HCl pH 8.0, 200mM NaCl, 1mM EDTA, 0.5mM EGTA, and protease inhibitors), and rotated at room temperature for 10 minutes, followed by centrifugation (1500 rpm at 4°C for 3 minutes). Supernatant was discarded and nuclei were resuspended/lysed in 1mL cold Lysis Buffer 3 (10mM Tris-HCl, pH 8.0, 100mM NaCl, 1mM EDTA, 0.5mM EGTA, 0.1% Na-Deoxycholate, and protease inhibitors) and transferred to pre-chilled 1mL Covaris AFA tubes (Covaris). Samples were sonicated using a Covaris S220 sonicator (High Cell Chromatin Shearing: PIP 140W, CPB 200, Duty Factor 5%, 15 minutes). Lysates were transferred to tubes and Triton X-100 was added (final 1%) followed by centrifugation (top speed, 10 minutes at 4°C in microcentrifuge). Supernatant was transferred to a new tube; protein concentration was measured by Bio-Rad Protein Assay Kit (Bio-Rad, Protein Assay Kit #5000002). Antibody-conjugated beads were washed 3 times in blocking buffer, resuspended in 50μL blocking buffer and added to 500μg input protein for overnight incubation with rotation at 4°C. 50μg lysate was stored at −20°C for input. On day 2, beads were washed 5 times in 1mL RIPA buffer (50mM HEPES-KOH pH 7.5, 500mM LiCl, 1mM EDTA, 1% NP-40, 0.7% Na-Deoxycholate) with inversion to mix followed by 2-minute incubation on ice for each wash. Beads were washed in 1mL final wash buffer (1xTE, 50mM NaCl) for 2 minutes at room temperature before final resuspension in 210μL elution buffer (50mM Tris-HCl pH 8.0, 10mM EDTA, 1% SDS). To elute, beads were incubated with agitation at 65°C for 30 minutes. 200μL eluate was removed to a fresh tube, and all samples (ChIP and reserved inputs) were reverse-crosslinked overnight at 65°C with agitation for a minimum of 12 hours, but not more than 18 hours. 200μL 1xTE was added to reverse crosslinked DNA to dilute SDS, and samples were treated with RNaseA (final 0.2mg/mL RNase; 37°C for 2 hours) and Proteinase K (final 0.2mg/mL Proteinase K; 55°C for 2 hours) before phenol:chloroform extraction and resuspension in 10mM Tris-HCl pH 8.0. ChIP and input DNA were quantified by Qubit (ThermoFisher) before library preparation using the NEBNext Ultra II DNA library prep kit according to manufacturer’s instructions (NEB). Samples were indexed for multiplex sequencing. Library quality was analyzed by BioAnalyzer (Agilent Genomics) and quantified using qPCR (NEB). Libraries were pooled for multiplex sequencing, re-quantified, and sequenced on the Illumina NextSeq500 or 1000 platforms (120 bp single-end sequencing; high output; Illumina).

#### Subcellular fractionation and western blotting

Cells were trypsinized and resuspended in fresh culture medium, then washed once in Dulbecco’s PBS. Cells were pelleted by centrifugation at 1,200*g* for 5 min at 4 °C. The pellet was then either processed with the Subcellular Protein Fractionation Kit for Cultured Cells kit (Thermo Fisher Scientific, 78840) or resuspended in 1× RIPA buffer (50 mM Tris-HCl pH 8, 150 mM NaCl, 0.1% SDS, 0.5% Sodium deoxycholate, 1% Triton X-100) with protease inhibitors to extract whole-cell lysate.

For subcellular fractionation, the manufacturer’s instructions were followed. For whole-cell lysate, the sample was nutated at 4 °C for 30 min and then centrifuged at 16,000*g* for 20 min at 4 °C. Supernatant containing protein was extracted to a new tube. Subcellular fractions and whole cell lysate were quantified using the Pierce BCA Protein Assay Kit (Thermo Fisher Scientific, 23225). Protein was stored at −80 °C until needed.

For blotting, protein was mixed with NuPAGE LDS sample buffer and sample reducing reagent (Thermo Fisher Scientific), denatured at 70 °C for 10 min and then cooled on ice. Then, 10–25 μg of sample was run on Mini-PROTEAN TGX Stain-free precast gels. The samples were transferred to a 0.2 μm nitrocellulose membranes for 1 h at 100V on ice. Membranes were blocked for 1 hour at room temperature in 5% milk in TBST, then incubated with primary antibodies diluted in 5% milk overnight at 4 °C. The following primary antibody concentrations were used: H3K9me3 (Abcam, ab8898, 1:1,000), H3K27me3 (Cell Signaling Technology, #9733, 1:1,000), H3 (Abcam, ab1791, 1:10,000), Cofilin (Cell Signaling Technology, 5175, 1:1000), GAPDH (Cell Signaling Technology, 5174, 1:1,000), hnRNPK (Santa Cruz, sc-28380, 1:1,000), Lamin B1 (Abcam, ab16048, 1:1,000), Lamin A/C (Santa Cruz, sc376248, 1:1,000), Lamin B Receptor (abcam, ab122919, 1:1,000). The next day, membranes were washed three times in TBST for 10 min each. The membranes were then incubated with secondary antibodies diluted in 5% milk TBST in the dark at room temperature for 1 h. The following secondary antibody concentrations were used: anti-rabbit HRP-linked (Cell Signaling Technology, #7074, 1:10,000), anti-mouse HRP-linked (Cell Signaling Technology, 7076, 1:10,000), anti-Mouse IRDye 800CW (LiCOR, 926-32210, 1:3,333), anti-rabbit Cy3 (Jackson ImmunoResearch Labs, 111-165-003, 1:6,666). The membranes were then washed three times in TBST for 10 min each and once in TBS for 5 min (all in the dark). Blots with HRP-linked secondary antibodies were additionally incubated for 4minutes with Clarity or Clarity Max ECL Western Blotting substrates (Bio-Rad). The blots were then imaged on the ChemiDoc MP Imaging System and analyzed using Bio-Rad Image Lab software.

#### Proximity labelling by TurboID

Plasmid vectors with the CMV promoter were designed to express either Lamin B1 fused to BirA and V5 tag or BirA-V5 alone. The TurboID protocol was performed according to the published protocol^59^ with some modifications. Cells were seeded in six-well plates with 2 ml of medium containing 4.5 × 10^5^ cells per ml. The next day, the medium was aspirated from each well and replaced with 2 ml of fresh medium before transfecting with a mixture of 5 μg of plasmid vectors, 250 μl Opti-MEM reduced serum medium (Thermo Fisher Scientific), 7.5 μl Lipofectamine 3000 reagent and 10 μl of P3000 (Thermo Fisher Scientific) reagent. After 48 h, the medium was aspirated from each well and replaced with medium containing 500 μM of biotin to enable proximity labelling. After 70 minutes, the plate was moved to ice, the medium was aspirated, and the wells were washed five times with 500 μl ice-cold PBS. Cells were pelleted at 300*g* at 4 °C for 3 min, the supernatant was aspirated, and the cell pellet was flash frozen in liquid nitrogen. Pellets were stored at −80°C.

Frozen cell pellets were resuspended in 5 pellet volumes Buffer A (10 mM HEPES pH 7.9, 5 mM MgCl2, 0.25 M sucrose+ 0.2 mM PMSF + 1mM DTT) and incubated on ice for 10 minutes (agitated halfway through incubation). Samples were centrifuged at 6,000*g* at 4 °C for 10 minutes, and the supernatant (cytosolic fraction) was transferred to a new tube for storage at −80°C. Pellets were then resuspended in 4 pellet volumes of Buffer B (10 mM HEPES pH 7.9, 1.5 mM MgCl2, 0.1 mM EDTA and 25% glycerol + 0.2 mM PMSF + 1mM DTT) with 0.42M NaCl and incubated on ice for 20 minutes with agitation. Samples were centrifuged at 9,400*g* at 4 °C for 15 minutes, and the supernatant (soluble nuclear fraction) was transferred to a new tube for storage at −80°C. Pellets were then resuspended in 4 pellet volumes of Buffer B (10 mM HEPES pH 7.9, 1.5 mM MgCl2, 0.1 mM EDTA and 25% glycerol + 0.2 mM PMSF + 1mM DTT) with 1M NaCl and incubated on ice for 20 minutes. Volume was brought up to approximately 450μl for efficient sonication. Samples were sonicated for 6-8 seconds on setting 4 twice, with 1 minute incubation on ice in between bursts (Dismembrator Model 100, Fisher). Samples were centrifuged at 9,400*g* at 4 °C for 15 minutes, and the supernatant (chromatin-bound fraction) was transferred to a new tube for storage at −80°C. The protein concentration was quantified using the Bio-Rad Protein Assay Kit (Bio-Rad, Protein Assay Kit #5000002).

To perform the immunoprecipitation (IP), 50 μl of MyOne Streptavidin C1 magnetic beads (Thermo Fisher Scientific) were washed twice in 1mL RIPA buffer before being combined with 75 μg of chromatin-bound lysate in a total volume of 500 μl of RIPA buffer (50 mM Tris-HCl pH 8, 150 mM NaCl, 0.1% SDS, 0.5% Sodium deoxycholate, 1% Triton X-100 + protease inhibitors) and incubated overnight in a rotator at 4 °C. The next day, the beads were collected on a magnet and washed and incubated in RIPA buffer twice for 2 minutes each time, 1 M KCl for 2 minutes, 0.1 M Na_2_CO_3_ for 10 seconds (without resuspension), 2 M urea for 10 seconds (without resuspension) and twice more in RIPA buffer for 2 minutes each time. The beads were then incubated at 95 °C for 15 minutes with 2 mM biotin, NuPAGE LDS sample buffer and sample reducing reagent (Thermo Fisher Scientific), and the supernatant was collected for Western blotting.

#### Cloning

Plasmids containing codon-optimized Lamin B1 driven by the CMV promoter were ordered through Twist Bioscience. For v5-BirA-Lamin B1 construct, the V5 epitope tag and the mini-Turbo BirA sequence (based off of the Addgene Plasmid #107176) were cloned into the N-terminus of Lamin B1.

#### RNA-seq

1.5 million HCT-116 cells were harvested after control or *HNRNPK* knockdown as described above. 4 biological replicates of each condition were collected for RNA-seq. Cells were washed with 1x Ca/Mg-free PBS and centrifuged at 100*g* for 15 minutes. Supernatant was aspirated off and the cell pellets were flash frozen and stored at −80°C. Cell pellets were shipped on dry ice to MedGenome. MedGenome extracted RNA and prepared libraries with Illumina Tru-Seq mRNA stranded kit. Libraries were sequenced on a Nova-seq with 100-bp paired end reads to a depth of at least 20 million reads.

## Quantification and statistical analysis

### HiDRO image analysis

Images were processed and analyzed as described previously^47^ with minor adjustments. Specifically, images from HiDRO plates were segmented and measured using CellProfiler^76^ (v.4.2.4). The edges of nuclei were detected using two-class Otsu thresholding^77^ and the edges of FISH spots were detected using three-class Otsu thresholding^77^. For spot segmentation, only spots within segmented nuclei were retained for measurement. Euclidean distance to the periphery was determined by calculating the distance from the centroid of each FISH spot to the nearest edge of corresponding nucleus as determined by the segmented DAPI edge. Normalized distances were determined by dividing the Euclidean distance by the nuclear area for each allele. The parameters for Otsu thresholding were tuned for each plate by visually inspecting the segmentation results in CellProfiler. To compare HiDRO measurements across different plates and experiments, robust *z*-scores for each measurement were generated by comparing each individual well against all of the wells on the same replicate plate. Negative control wells with non-targeting siRNA were included for comparison. The mean of each measurement was calculated per well, and the formula for robust *z*-score was applied: (mean measurement for well − median of mean measurements for all wells on plate excluding positive controls)/(1.486 × (median absolute deviation for the mean measurements across the plate, excluding positive controls)). Full data are reported in Supplementary Table 2. For the validation de-pooled screen, robust *z*-scores were calculated from a null distribution of at least 50 wells seeded with non-targeting control siRNA 5. Full data are reported in Supplementary Table 4. To correct for chromatic aberrations, we recorded the chromatic offset in x,y (for 2D analysis of HiDRO images) with 0.1 µm TetraSpeck microspheres (Thermo Fisher Scientific, T7279). Microscope calibrations were then performed through mechanical and software modifications to correct for the chromatic offset during subsequent acquisitions using MetaXpress software. We continually took measurements with Tetraspeck beads and recalibrated our system before each experimental run if we recorded an offset greater than the size of a pixel of that system. Note that we cannot completely rule out that some chromatic shifts transiently occurred throughout our long HiDRO imaging sessions (24 h per plate × 6 plates per week). Thus, to account for additional plate-to-plate variation, we prepared replicate plates on separate weeks and restricted our analysis of screen results to robust *z*-scores (Supplementary Table 2) rather than the absolute values of Euclidean distance to the periphery defined by the DAPI edge.

### Primary screen and validation screen hit selection

Hits were selected from the primary screen data by meeting the following criteria based on nuclei count (≥ 100 nuclei across both replicates) and Euclidean and normalized distance to the periphery measurements for the LAD and nonLAD probe: both replicates had *z*-scores with a value of ≥1.85 for the Chr2q-LAD’s Euclidean distance to the periphery. Both replicates had *z*-scores with a value of ≥1.5 for the Chr2q-LAD’s normalized distance to the periphery. Both replicates had *z*-scores with a value ≤ 1.5 for the Chr22q-nonLAD’s Euclidean and normalized distance to the periphery. Finally, the gene had to be expressed in HCT116 cells (>50 DESeq2 normalized counts from control HCT116 cells). These cutoffs determined the 101 primary screen hits. In the primary screen, each gene was targeted with a pool of four siRNA duplexes in a single well.

For our validation screen, we ordered from ICCB-L 384-well plates seeded with the four siRNA duplexes for each primary gene hit de-pooled into four separate wells, arranged randomly on one 384-well plate. We performed the validation screen in triplicate. Robust *z*-scores for overlap measurements were calculated for each well versus a null distribution of at least 50 wells (Human 6&7 or Human 8) seeded with non-targeting control siRNA 5 (Supplementary Table 6). We set a nucleus count cut-off of 50 nuclei. Individual siRNA duplexes were considered to be validated hits if they had *z*-scores with values ≥ 1.85 in at least 2 out of 3 replicates for the Chr2q-LAD’s Euclidean distance to the periphery and *z*-scores with values ≥ 1.5 in at least 2 out of 3 replicates for the Chr2q-LAD’s normalized distance to the periphery. This resulted in 117 out of 208 duplexes validating corresponding to 51 out of 52 primary hit genes. This left us with 51 out of 52 validated genes, which are listed in Supplementary Table 4. Genes validating with or 2 more duplexes were considered high-confidence hits.

### DNA and RNA slide FISH image analysis

Widefield images were deconvolved with Huygens Essential v.20.04 (Scientific Volume Imaging) using the classic maximum-likelihood estimation algorithm with theoretical PSFs. Additional settings include signal-to-noise ratio (SNR) of 40 and 50 iterations for FISH spots or SNR of 40 and 2 iterations for DNA stain. The deconvolved images were segmented and measured using a modified version of the TANGO 3D-segmentation plug-in for ImageJ ^78^. Nuclei and FISH spots were segmented using a hysteresis-based algorithm. To correct for chromatic aberrations, we recorded the chromatic offset in x,y,z with 0.1 µm TetraSpeck microspheres (Thermo Fisher Scientific, T7279). Microscope calibrations were then performed through mechanical and software modifications to correct for the chromatic offset during subsequent acquisitions using Leica software. Remaining corrections were applied to images immediately after acquisition.

### RNA-seq analysis

Raw fastq reads were aligned to the hg38 reference genome with RefSeq transcripts aligned by UCSC (https://hgdownload.soe.ucsc.edu/goldenPath/hg38/bigZips/genes/) using STAR^79^ (v2.7.1a) with parameters: --outFilterMismatchNmax 3 --outFilterMultimapNmax 1. Gene-level raw read counts were obtained using featureCounts^80^ (subread v2.0.6) with parameters: -t exon -g gene_id -p --countReadPairs -B -Q 20. Genes with zero counts across all 8 samples were excluded from downstream analyses. Raw counts were normalized to Transcripts Per Million (TPM) using gene length obtained from featureCounts. Then, differentially expressed genes were called using DESeq2^81^ (v1.46.0). Batch effects were controlled by adding batch factor to the DESeq2 design (design = ∼ batch + condition). Log_2_ fold change was shrinked using the apeglm method^82^.

Calculation of gene density within LADs: The number of genes in each LAD was calculated by overlapping LAD coordinates with gene promoters (promoters defined as 1500 bp upstream of TSS, 500 bp downstream). The non-LAD control regions were created by shuffling LADs using bedtools^83^ (v2.31.0) with parameter: -excl -chrom. 100 shuffles were performed. For each shuffle, the number of genes overlapping each non-LAD control region was counted. For each LAD region, the median of the 100 shuffled regions was used to represent corresponding non-LAD control.

Gene ontology analysis: For genes overlapping each LAD category, gene ontology analysis was performed using clusterProfile^84^ (v4.14.4) with parameters: ont = “BP”, pAdjustMethod = “BH”, pvalueCutoff = 0.05, qvalueCutoff = 0.05. The top 15 biological processes were selected if more than 15 terms were significantly enriched.

Distribution of differentially expressed genes within LADs: Each LAD region was divided into 100 bins (bin 1 – 100), and genes were assigned to a bin based on their transcription start sites. Genes located −100 kb and +100 kb of 5’ and 3’ end of LADs were assigned to bins −9-0 and 101-110 respectively. If a gene was adjacent to multiple LADs, it was assigned to the closest LAD.

### ChIP-seq analysis

ChIP-seq libraries were sequenced on an Illumina NextSeq 500/1000 platform (Illumina, San Diego, CA) using single-end 120 bp reads to a target depth of at least 26 million uniquely mapped reads per library (Supplementary Table 7). Base call (BCL) files were demultiplexed in Illumina BaseSpace Sequence Hub using the sample-specific dual-index combinations provided during library preparation. Demultiplexing was performed with the integrated bcl2fastq pipeline, and the resulting FASTQ files were downloaded for downstream analysis.

Read quality was assessed using FastQC^85^ (v0.12.1) and summarized with MultiQC^86^(v1.14). Adapter trimming, low-complexity read removal, and PCR duplicate removal were performed with fastp^87^ (v0.23.2) using default parameters unless otherwise stated. Processed reads were aligned to the human reference genome (GRCh38) using Bowtie2^88^ (v2.5.1) with default parameters. Reads with mapping quality (MAPQ) ≤ 20, unmapped reads, secondary alignments, or PCR duplicates were filtered out using Sambamba^89^ (v1.0.0) with the expression: ‘mapping_quality>=20 and ref_name =∼ /^chr[0-9XY]/ and not duplicat’.

To ensure equal sequencing depth across libraries and to mitigate coverage biases, filtered BAM files were randomly downsampled using samtools^90^ (v1.18) with the ‘view -b -s’ option to match the library with the smallest number of reads per condition, resulting in ≥ 30 million uniquely mapped reads per library. Biological replicate concordance was assessed using deepTools ‘plotCorrelation’^91^ (v3.5.1) with Spearman correlation coefficients calculated in 50 kb bins. Replicates with high correlation coefficients were merged using SAMtools ‘mergè for downstream analysis.

#### Coverage track generation, and domain calling

Coverage tracks for Lamin B1, H3K9me2, H3K9me3, and H3K27me3 ChIP-seq datasets were generated by extending reads to the estimated fragment length (120 bp) and normalizing to counts per million (CPM). The enrichment of the IP libraries over their corresponding inputs was calculated as a log₂(IP/Input) signal in 1 kb bins genome-wide using deepTools bamCoverage and bamCompare. The resulting signal was smoothed with a 10 kb sliding window and *z*-score normalized per chromosome using a custom awk script. Regions present in the ENCODE blacklist for hg38^92^ were excluded from all analyses. Normalized coverage tracks were visualized using pyGenomeTracks^93,94^ (v3.8).

Domains of enriched ChIP-seq signal were identified using a 2-state Hidden-Markov Model implemented in HMMDomainCaller^95^ with minor modifications (https://github.com/rikrdo89/HMM_caller). The HMM was trained on ChIP-seq signal from control samples, using a bin size of 5kb. Following HMM domain calling, we applied a post-filtering step in which domains were discarded if either their length or their ChIP-seq enrichment score fell below the 25th percentile of all called domains in the control condition. LAD calls were compared to previously published datasets using BEDTools^83^ (v2.31.1), BEDOPS^96^ (v2.4.41), eulerR (v7.0-0), and Intervene^97^ to assess concordance.

#### LAD Categorization and ChIP-seq Signal Enrichment

LADs identified in control cells were compared to LADs from hnRNPK-depleted (siHNRNPK) cells using BEDOPS ‘bedmap’ with the ‘--echo --bases --bases-uniq-f’ options to calculate fractional base-pair overlap^96^. Control LADs were classified as: i) Retained: ≥ 75% overlap, ii) Weakened: 25–75% overlap, iii) Lost: ≤ 25% overlap. These thresholds were supported by inspection of genome browser tracks and heatmaps of Lamin B1 enrichment over called domains.

Mean normalized ChIP-seq signal over LAD categories was computed with deepTools ‘computeMatrix scale-regions’ and visualized using ‘plotProfilè. For genome-wide comparisons, coverage matrices were generated by scaling all LADs to their median size. ChIP-seq signal distributions between control and siHNRNPK conditions were compared using dsCompareCurves (v1.2.0) with paired Wilcoxon signed-rank tests (two-sided, p < 0.05). Statistical robustness was assessed by generating 1000 bootstrap resamples of the paired differences to estimate 95% confidence intervals. Regions passing the Wilcoxon significance cutoff were flagged as differentially occupied. Mean difference curves and confidence intervals were plotted in R (v4.3.2) using ggplot2 (v3.4.4).

### Statistics and reproducibility

The number of samples per condition (*n*) is indicated in the respective figure legends of all graphs. For HiDRO experiments, each well was a biological replicate. Each 3D slide FISH experiment reported in this study was repeated in at least one biological replicate (Fig. 2c, Supp. Fig. 2c, Fig. 5e-g, Supp. Fig. 5k). Immunofluorescence experiments were repeated in at least one biological replicate (Fig. 2f, Supp. Fig. 5b). The Lamin B1 Turbo-ID experiment in Fig. 2d was repeated in one additional biological replicate (Supp. Fig. 2e). Western blot experiments were repeated in at least one additional biological replicate (Fig. 2e, Supp. 2b,g, Supp. Fig. 5a). Protein class designations were determined by PANTHER ^50^ and manual curation of unassigned proteins. All other statistical tests are discussed in the corresponding methods section or figure legend. Statistical analyses were performed using either R (v.4.4.1), Python or Prism 10 software (GraphPad; v.10.4.1).

## Data availability

Datasets reported in this paper are available at the Gene Expression Omnibus under accession number GSE303855, which will be made public following acceptance.

## Code availability

CellProfiler pipeline for image segmentation is available at Zenodo (https://doi.org/10.5281/zenodo.7699078).

